# Haplotype-resolved genome architecture mapping uncovers pervasive structural heterogeneity between human homologous chromosomes

**DOI:** 10.64898/2026.06.16.732681

**Authors:** Julia Markowski, Alexander Kukalev, Claudia Robens, Christoph J. Thieme, Adam Streck, Hui Mao, Krishna Mohan Parsi, René Maehr, Ana Pombo, Roland F. Schwarz

## Abstract

Resolving the three-dimensional structure of chromatin with haplotype specificity remains a fundamental challenge for understanding genetic and epigenetic contributions to gene regulation in healthy development and in disease. Phasing of chromatin contacts derived by ligation-dependent methods relies on high single-nucleotide variant (SNV) density, which limits its applicability in human genomes where SNVs are sparse. Here, we present CoPhasing, a strategy that leverages the intrinsic haplotype fidelity of Genome Architecture Mapping (GAM), a ligation-free method that inherently captures haplotype-matched genomic neighborhoods in thin nuclear slices. CoPhasing assigns SNV-free reads to their haplotype through proximity to informative reads, enabling efficient and accurate phasing even in SNV-sparse regions. Applying CoPhasing genome-wide reveals extensive, previously inaccessible structural heterogeneity between human homologous chromosomes, demonstrating the power of GAM CoPhasing to investigate allele-specific differences in 3D genome organization and their relevance to gene regulation and disease.

## Introduction

Three-dimensional genome architecture is fundamental to the regulation of gene expression, through enhancer-promoter interactions, chromatin loops, topologically associating domains (TADs), active and repressive chromatin compartments, and ultimately chromosome territories in the nucleus. Increasing interest in understanding the impact of spatial genome interactions has led to significant advances in both imaging and sequencing-based, experimental and computational methods^1^. Not only have these methods helped to uncover novel concepts of nuclear organization^2,3^, they have also enabled a mechanistic understanding of gene expression deregulation, e.g. through unravelling the effects of chromatin architecture in various diseases^4–6^.

Although previous studies provided important insights into 3D chromatin organization, they largely treated the genome as a single entity, disregarding differences in 3D structure between homologous chromosome pairs. Distinct 3D structures between the homologous chromosomes have been observed across multiple systems, including in *C. elegans*^7^, in healthy mouse and human cells^8–11^ and in cancer^12^. Resolving haplotype-specific chromatin conformation is therefore key for an improved understanding of the haplotype-specific regulatory landscapes of genomes, and their impact on health and disease.

In sequencing-based approaches, identifying haplotype-specific differences first requires an assembly of heterozygous germline SNVs into parental haplotypes, followed by phasing of sequencing reads to their haplotype of origin^13^. Current phasing methods generally assign sequencing reads to their respective haplotype only when those reads directly overlap heterozygous SNVs - an approach we refer to as *direct phasing*. However, homologous chromosomes differ in sequence only at few heterozygous variant sites. For example, in the human genome, heterozygous SNVs are observed every 1-1.5 kb^14^. The sparsity of genetic variants poses a challenge to studying the individual 3D structure of homologous chromosomes. Previous attempts to investigate haplotype-specific chromatin contacts in human Hi-C data through direct phasing reported read phasing efficiencies of only ∼3%^13^, ∼7.5%^15^, up to ∼11%^16^, depending on the SNV density available. Those approaches prove inefficient, as they discard the majority of reads and thus lead to substantial information loss, severely limiting haplotype-specific chromatin conformation analyses.

Genome Architecture Mapping (GAM)^10,17^ provides the unique opportunity to address this challenge. GAM infers the nuclear organization of chromatin by identifying spatially close DNA through their increased co-segregation in hundreds of thin cryosectioned nuclear slices (called nuclear profiles, NPs). Calculating the frequency of co-observations of pairs of genomic loci across all NPs enables the construction of chromatin contact maps^17^. Due to the physical sampling process based on thin nuclear slices, recurrently co-isolated DNA fragments most likely originate from the same chromosomal copy^17^, enabling the reconstruction of chromosome-spanning haplotypes with high accuracy^18^.

Here, we introduce CoPhasing, an approach that exploits the inherent haplotype fidelity of the sampling process of GAM to enable phasing of the large majority of sequencing reads, including those not overlapping heterozygous SNVs. By overcoming the fundamental limitation of direct phasing, CoPhasing generates high-quality haplotype-specific chromatin contact matrices in high-resolution, irrespective of the SNV density of the underlying genome. Applying CoPhasing to GAM data of human cells, we detect extensive and previously unappreciated differences in haplotype-specific chromosome structure across the human genome, opening new opportunities for the genome-wide, haplotype-resolved study of 3D chromatin organization in development and disease.

## Results

### Limitations of direct phasing in SNV-sparse human genomes

To evaluate the impact of SNV density on the generation of haplotype-phased chromatin contact matrices from GAM data, we used direct phasing on GAM datasets from a SNV-dense and a SNV-sparse genomic background.

First, we replicated our previous work in the hybrid mouse embryonic stem cell (mESC) line F123^10^, a cross between the divergent S129 and CAST strains that is characterized by high SNV density (1 SNV every ∼132 bp). Direct phasing of the F123 genome generates informative phased chromatin contact matrices that revealed extensive folding variability between homologous chromosomes^10^ (**Fig. 1a**).

**Fig. 1:**
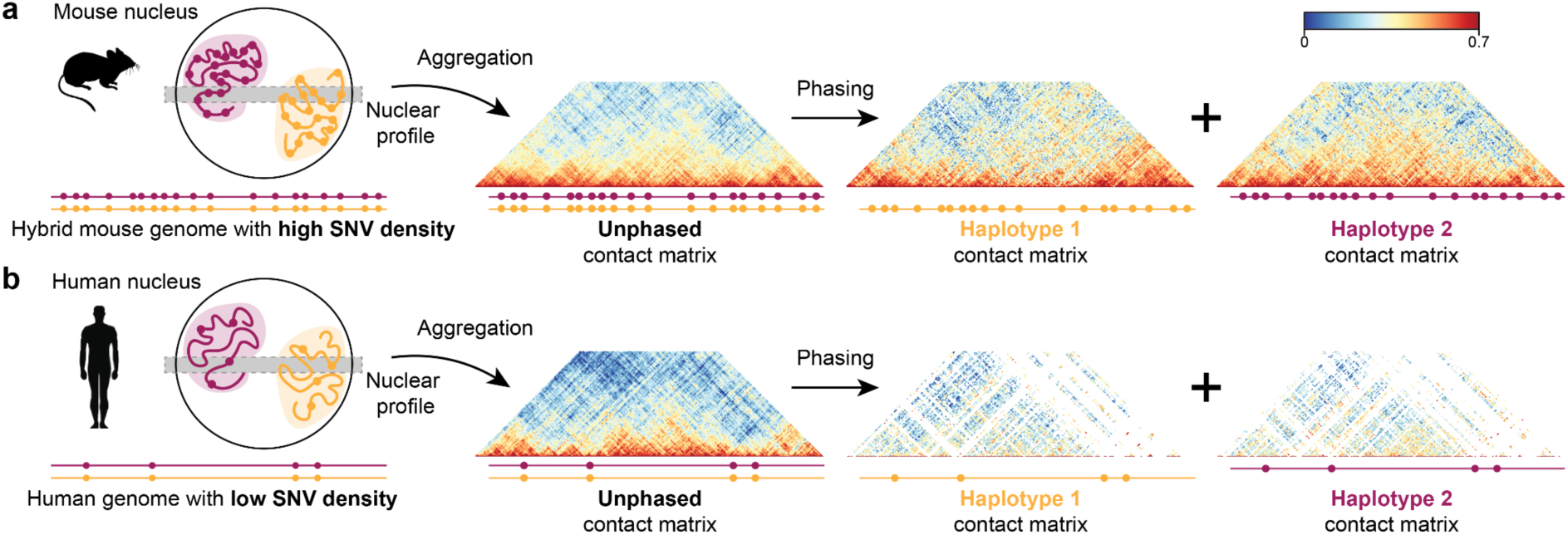
Low SNV density in human genomes limits the quality of phased GAM contact matrices. **a**, Schematic showing the chromatin folding of homologous regions (yellow and red) in a nucleus of a hybrid mouse with high SNV density. A thin GAM nuclear profile (gray bar) captures DNA fragments in 3D space. Aggregation of these captured fragments across multiple profiles enables the reconstruction of saturated unphased contract matrices. The high SNV density of the hybrid mouse genome yields enough SNV-containing reads which can be assigned to their haplotypes, resulting in high-quality phased contact matrices (example region: chr7: 48 Mb - 58 Mb). **b,** In human genomes, markedly lower SNV density yields fewer haplotype-informative and thus phasable reads. While unphased GAM contact matrices are well-saturated (chr7: 48 Mb - 58 Mb), the scarcity of SNVs leads to an insufficient haplotype assignment of reads and ultimately to sparse phased contact matrices.

Next, we used direct phasing on a GAM dataset of the human embryonic stem cell (hESC) line H1^3^, a well-characterized, haplotype-resolved^8^, cell line widely used in the research of 3D genome structure^3^. Its markedly lower SNV density of 1 SNV every ∼2 kb yielded only ∼4% of phased reads, consistent with the limitations described for ligation-based assays. While the unphased H1 hESC contact matrix is well saturated and informative, the directly phased matrices are sparse and insufficient for haplotype-specific chromatin conformation analyses (**Fig. 1b**), confirming that the low SNV density of the human genome poses a fundamental barrier to direct phasing approaches.

### Design and implementation of the CoPhasing approach

We designed CoPhasing based on key principles of chromosome organization and the process of spatial genome sampling in GAM technology. First, the chromosomes of mammalian cells typically occupy discrete chromosome territories in the nucleus, with homologous chromosomes often located far from each other^19^. Through the ultra-thin slicing process of GAM, each nuclear profile (NP) contains only ∼5% of the genome, with minimal chance of the same homologous region from both chromosome copies being captured in the same nuclear slice. Orthogonal imaging analyses by fluorescence in situ hybridization shows that homologous 30kb genomic regions co-occur in less than 1% of NPs^20^.

Second, because each DNA fragment captured in a single NP originates from one of the two homologous chromosomes, all sequencing reads derived from that DNA fragment must originate from the same haplotype (**Fig. 2a**, red haplotype). Due to GAM’s inherent haplotype fidelity^18^, even fragments in close proximity to each other on the linear genome are highly likely to have originated from the same homologous chromosome (**Fig. 2a**, yellow haplotype). We therefore hypothesised that the inherent properties of spatial genome sampling in GAM could be leveraged to reliably assign the haplotype of SNV-lacking sequencing reads by co-phasing them to the nearest phased SNV detected in the same NP.

**Fig. 2:**
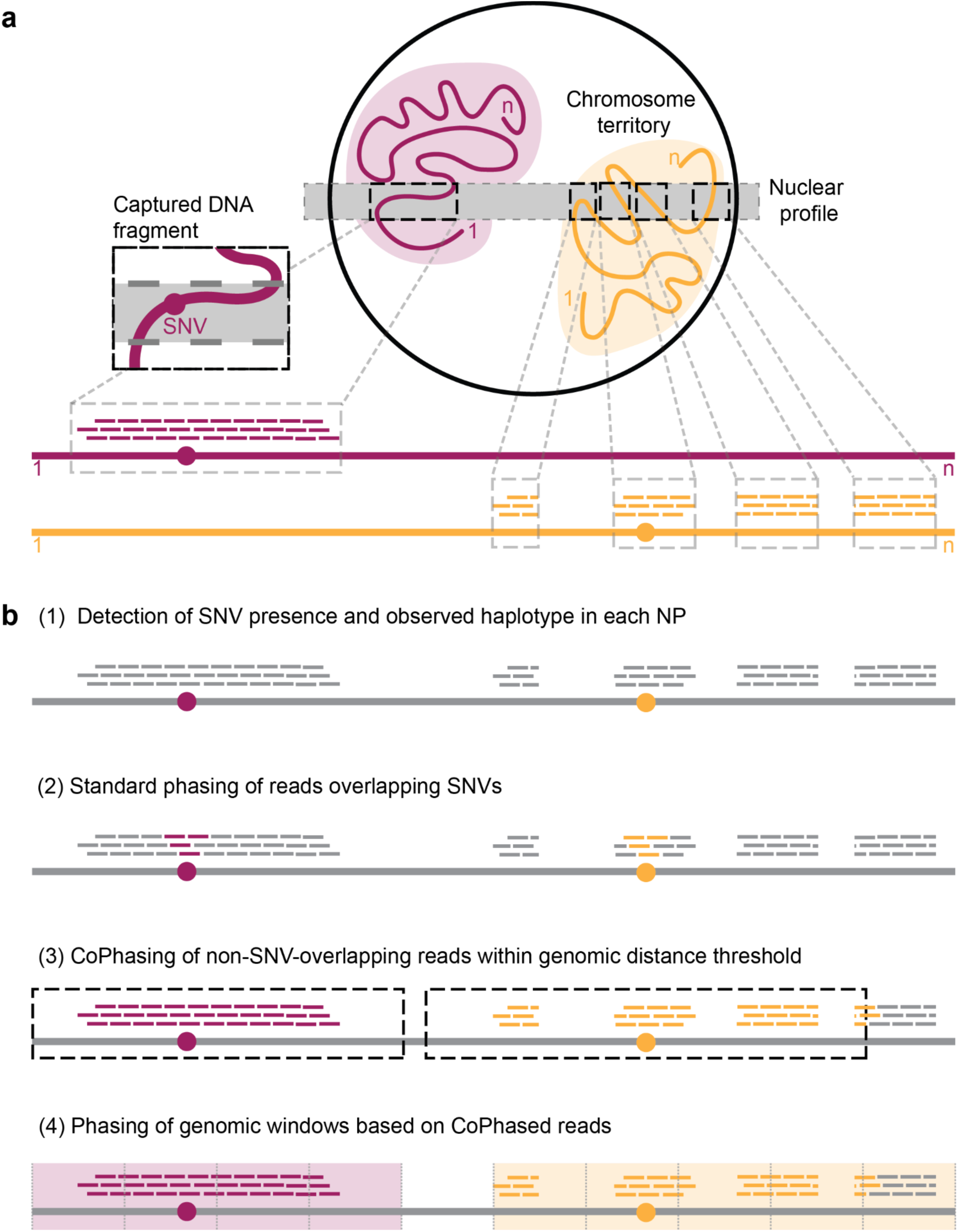
The CoPhasing concept of GAM sequencing data. **a**, Reads derived from continuous (red) and contiguous (yellow) DNA fragments captured in a nuclear profile originate from their corresponding homologous chromosomes and haplotypes. **b,** After alignment of all sequencing reads and identification of observed SNVs in each NP, SNV-overlapping reads are haplotyped. Reads not overlapping SNVs are assigned to the haplotype of their closest SNV-overlapping read within a genomic distance threshold (cophased), and genomic windows are phased accordingly.

In the first step of CoPhasing, the alleles of present SNVs are determined separately for each individual NP (**Fig. 2b(1), Fig. S1**). Then, sequencing reads that directly overlap SNVs are assigned to their respective haplotype (**Fig. 2b(2)**). All reads that do not overlap SNV positions are identified within a defined genomic distance and assigned to the haplotype of their closest phased read, hereinafter referred to as “cophased” (**Fig. 2b(3)**). Finally, to construct chromatin contact matrices in high resolution, the genome is partitioned into contiguous, non-overlapping windows of constant size, which are assigned to a haplotype based on the phased coverage within that window, as previously described^10^ (**Fig. 2b(4)**).

### Benchmarking and parameter estimation in a hybrid mouse genome

Since the low SNV density of the human genome is the limiting factor in generating haplotype-specific chromatin contact matrices through direct phasing, there is no gold standard to benchmark CoPhasing results against. We therefore turned to F123 mESC as a benchmark system, downsampling its dense SNV set from 1 SNV per ∼132 bp to 1 SNV per ∼2 kb, to match the SNV density of H1 hESC (**Fig. S2a**). Using the downsampled F123 SNV set for CoPhasing of the F123 mESC GAM dataset allowed us to assess CoPhasing efficiency and accuracy at the read, window, and matrix levels under human-like conditions, while retaining the full F123 direct phasing results as ground truth (**Fig. 3a**).

**Fig. 3:**
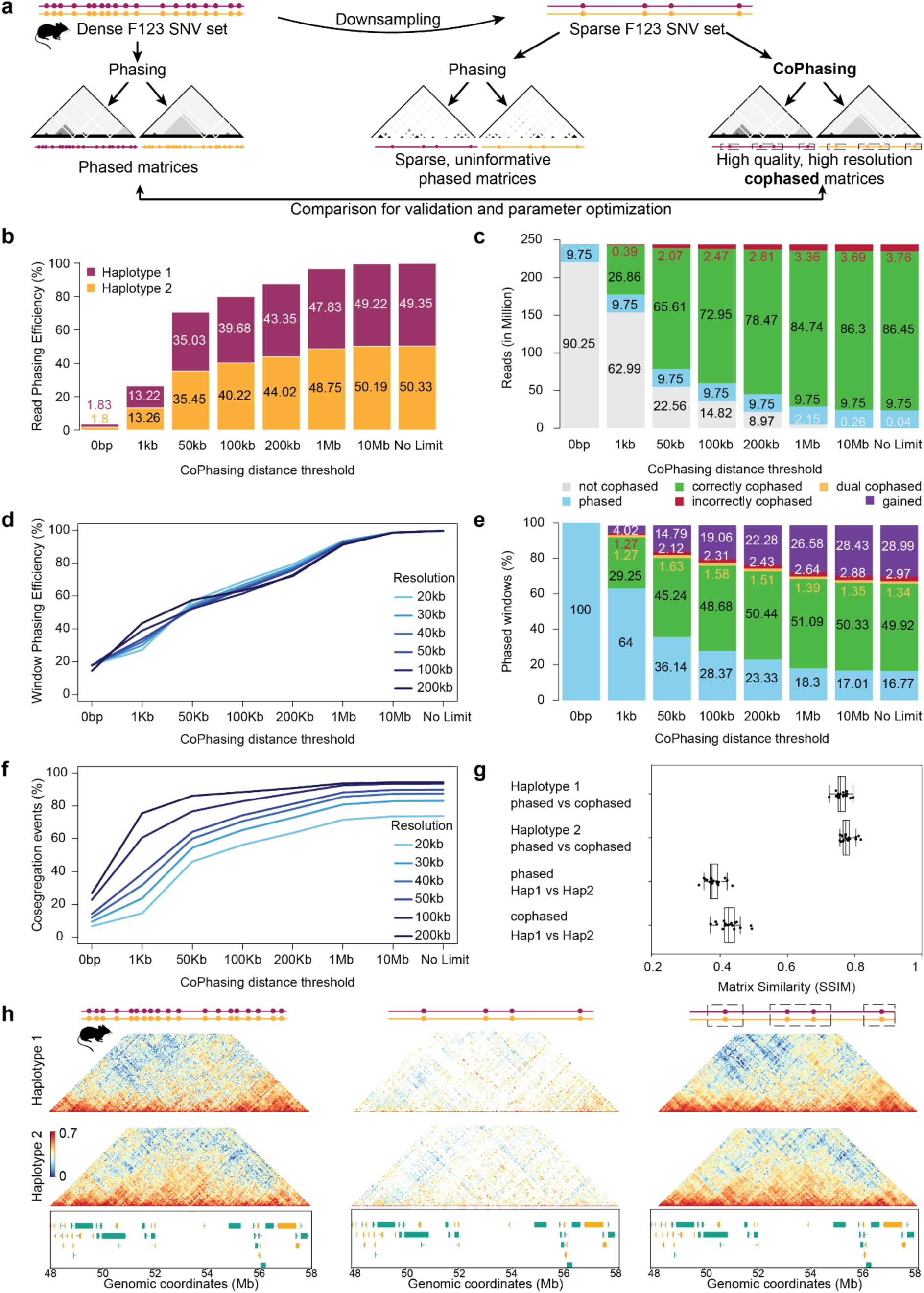
Benchmarking and parameter estimation across read-, window-, and matrix level. **a,** Dense F123 mESC SNV set was downsampled to human SNV density (see **Fig. S2**). Haplotype-specific contact matrices were generated from GAM data with direct phasing based on the full and downsampled SNV set as reference. CoPhasing results using the downsampled SNV set with human SNV density were compared to the results from direct phasing using the full SNV set to investigate phasing efficiency and accuracy across read-, window-, and matrix level. **b,** Read phasing efficiency increases with larger CoPhasing distance thresholds. **c,** Read phasing accuracy across increasing CoPhasing distance thresholds. Haplotype assignment of cophased reads is compared to phase-informative reads overlapping SNVs from the full F123 SNV set (∼250 million reads). Approximately 10% of reads overlapped SNVs in the downsampled SNV set (blue). **d,** Window phasing efficiency across increasing CoPhasing distance thresholds and resolutions. **e,** Window phasing accuracy across increasing CoPhasing distance thresholds. Purple fractions show windows recovered using CoPhasing, for which direct phasing provides no information and thus accuracy cannot be determined. **f,** Percent of non-zero cosegregation events across increasing CoPhasing distance thresholds and resolutions. **g,** Structural Similarity Index Measure (SSIM) comparing haplotype-specific chromatin contact matrices generated by direct phasing based on the full SNV set and CoPhasing on the downsampled SNV set. **h,** Haplotype-specific chromatin contact matrices derived from GAM data directly phased on the full F123 SNV set (left), directly phased on the subsampled F123 SNV set (center), and using CoPhasing on the subsampled F123 SNV set (right), on a 10 Mb genomic region (chr7: 48 Mb - 58 Mb).

Direct phasing of the GAM dataset with the full F123 SNV set discarded most reads, assigning only 37% of the total uniquely mapped sequencing reads to their haplotype of origin, as previously shown^10^. In contrast, direct phasing of the downsampled SNV set, resulted in phasing of fewer than 4% of all reads (**Fig. 3b**, 0 bp distance threshold), replicating the low phasing efficiency observed in H1 hESC, and confirming that the downsampled F123 mESC SNV set accurately models the SNV density constraints of the human genome.

To explore the effect of distance between each SNV-lacking read to the nearest SNV, we applied CoPhasing with different distance thresholds (**Fig. 3b**). CoPhasing within 1 kb, assigns 26% of reads, almost reaching the read phasing efficiency of direct phasing on the full F123 SNV set. At 50 kb, over 70% of reads were assigned to a haplotype, and at a threshold of 10 Mb, nearly all reads could be assigned (>99%; **Fig. 3b**). Cophased reads were balanced between the two haplotypes at all distance thresholds.

To assess the accuracy of CoPhasing and its dependency on CoPhasing distance, we compared the haplotype assignments of cophased reads to the assignment of the 37% subset of reads overlapping an SNV from the full F123 set (**Fig. 3c**). At 1 kb CoPhasing distance threshold, 27% of reads were cophased in addition to the 9.75% of reads directly phased, with only 0.4% assigned to the incorrect haplotype. The proportion of correctly cophased reads increased with larger CoPhasing distance thresholds while the rate of incorrect assignments remained below 3% at 200 kb distance, supporting the fact that GAM slices capture neighboring genomic regions from individual haplotypes^18^. Enabling cophasing to the maximum chromosome length without a distance threshold resulted in 96.2% reads correctly matched to their haplotype of origin and only 3.76% incorrectly assigned. Only 0.04% of reads could not be assigned to a haplotype due to absence of any SNV overlap in that chromosome and NP. Overall, CoPhasing shows exceptionally high read phasing efficiency and accuracy of haplotype assignment, even using long CoPhasing distances or no distance threshold, nominally reaching 195 Mb for the largest mouse chromosome.

To aggregate individual read-level information into structured GAM contact maps, the genome was binned into contiguous genomic windows of defined size, ranging from 20 kb to 200 kb for evaluation of different resolutions. Smaller windows offer finer, more detailed chromatin contact information in higher resolution, but their detection requires sufficient read coverage within each window. With direct phasing on the full SNV set, only 70 - 75% of genomic windows can be phased at resolutions of 10 - 100 kb^10^. CoPhasing enables phasing of nearly all genomic windows (>95.5%) stably across different resolutions at the low human SNV density using a 10 Mb CoPhasing distance threshold (**Fig. 3d**). The fraction of incorrectly assigned windows remained consistently below 3% across all CoPhasing distance thresholds (**Fig. 3e**). CoPhasing was even able to correctly assign rare dual phasing cases detected in the full-SNV maps, which occur when DNA from both haplotype copies is captured within a single GAM slice, also seen in direct phasing.

Chromatin contacts in GAM data are detected by quantifying how frequently pairs of genomic windows are co-observed across NPs, represented in cosegregation matrices normalized to window detection frequencies. The percent of cosegregating window pairs consistently increases up to 1 Mb with increasing CoPhasing distance thresholds, after which additional gains are minimal (**Fig. 3f**). A suitable resolution of the cophased contact matrices was determined as 20 kb by assessing the quality genome sampling, as described previously^10^.

Finally, we generated haplotype-specific F123 chromatin contact matrices at 20 kb resolution based on the subsampled SNV set with human SNV density, using a 10 Mb CoPhasing threshold. We then compared the cophased matrices to directly phased matrices based on the full and downsampled SNV set (**Fig. 3g,h**). We systematically assessed the similarity between phased matrices derived from direct phasing and CoPhasing by calculating their Structural Similarity Index Measure (SSIM). We found that both haplotypes show high SSIM values between matrices generated using CoPhasing on the reduced SNV set and direct phasing on the full SNV set, and that the differences between haplotypes were highly comparable between the two sets, as shown by highly similar SSIM values (**Fig. 3g)**. While direct phasing on the full SNV set yielded detailed, haplotype-specific contact matrices (**Fig. 3h** left), direct phasing on the downsampled SNV set resulted in extremely sparse haplotype-specific contact matrices (**Fig. 3h** middle). CoPhasing rescued the haplotype-specific chromatin contact information, and enabled the recreation of highly detailed haplotype-specific matrices from low SNV density that show remarkable similarity to the benchmark (**Fig. 3h** right). Thus, CoPhasing effectively restored haplotype-specific contact differences in low SNV density.

Overall, benchmarking in the F123 mESC line showed that CoPhasing reliably recovers haplotype-specific chromatin contact matrices and restores the otherwise inaccessible haplotype-specific chromatin interaction signal at low SNV density. Beyond the low SNV density context, applying CoPhasing on the full F123 SNV set further increased the efficiency of window phasing to 99.65% at 30 kb resolution compared to 73.72% obtained through direct phasing, demonstrating that CoPhasing improves GAM phasing performance even in SNV-dense genomes and should therefore be considered the default phasing approach for GAM data. The high-resolution cophased and unphased segregation tables generated from the F123 mESC GAM dataset with the full set of SNVs was made available in a repository^21^.

### High-resolution haplotype-specific chromatin conformation matrices in human reveal differential contacts

Before applying CoPhasing to human GAM data, we confirmed that after direct phasing each H1 hESC NP samples one single haplotype (98.4% ± 2.5%, mean ± SD), across all NPs and chromosomes, while mixed alleles are rarely detected (<3.3% ± 4.3%, mean ± SD). Having confirmed the haplotype fidelity in the H1 hESC GAM dataset, we applied CoPhasing across a range of distance thresholds to generate haplotype-specific chromatin contact matrices. Consistent with the benchmarking performance, CoPhasing of H1 hESC GAM data using a 10 Mb distance threshold enhanced the efficiency of read phasing from below 4% to over 96%, resulting in an increase in genomic window detection from below 13% to over 96%, and a detection of co-segregating windows pairs of over 80% at 20 kb resolution and 90% at 40 kb resolution (**Fig. S2b-d**). Visualization of contact matrices generated with different CoPhasing distance thresholds (0 bp up to full chromosome length) demonstrates the increased sampling of co-segregation events, and shows significant structural differences between H1 hESC haplotypes, irrespective of distance cutoff (Methods, **Fig. S3**). We chose a resolution of 40 kb for further analyses of the cophased pairwise contact matrices based on genome sampling quality across different resolutions and genomic distances (Methods, **Fig. S4**).

CoPhasing of H1 hESC GAM data achieves haplotype-resolved detection of chromatin interactions, revealing pronounced haplotype-specific chromatin conformation across multiple genomic scales. Clear differences in contact frequency emerge between cophased haplotypes, while unphased contact maps appear to represent a composite of both haplotypes (**Fig. 4a**). To identify statistically robust haplotype-specific differences in cophased H1 hESC chromatin contact maps, we applied a permutation test in which segregation tables for both haplotypes were pooled and randomly split into two permuted haplotypes, and empirical contact differences were compared to the randomized differences across 2000 replicates (Methods, **Fig. S5**). The statistical analysis using the permutation test revealed differential contacts showing stronger contact on haplotype 1 or haplotype 2, consistent with visual differences observed in cophased and unphased contact maps (**Fig. 4a**). We assessed the genome-wide magnitude of structural differences and identified a consistent fraction of on average ∼8% of chromatin contacts as differential between homologous chromosomes, balanced with ∼4% specific to each haplotype (**Fig. 4b, Fig. S7b**).

**Fig. 4:**
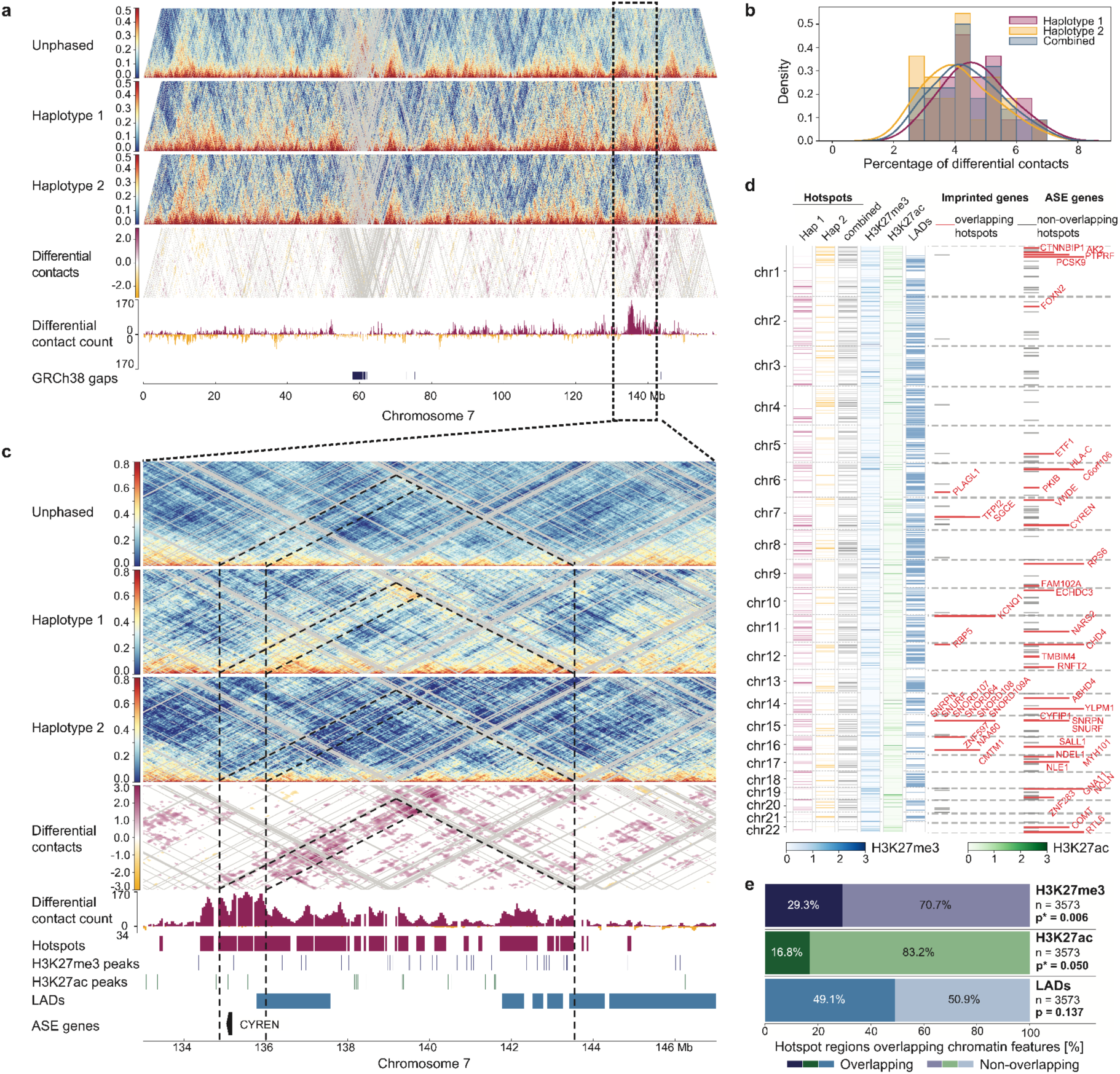
Differential contacts at ASE locus in H1 hESC and hotspot regions enriched for Polycomb revealed by CoPhasing. **a,** Unphased and cophased chromatin contact matrices, differential contacts as identified by permutation test (red: stronger contact on haplotype 1, white: no difference detected, yellow: stronger contact on haplotype 2) in chromosome 7. CoPhasing-derived matrices of H1 hESC generated using 40 kb resolution and 10 Mb CoPhasing distance threshold. Red indicates frequent interactions between respective genomic regions and blue no interactions. Differential contact count based on permutation test results are visualised per haplotype (haplotype 1: red, haplotype 2: yellow). Filtered regions in the matrices are depicted in light grey and regions with low mappability in GRCh38 genome are indicated below (mean mappability ≤ 0.1). **b,** Density distribution of percentage of differential contacts in the H1 hESC genome across chromosomes per haplotype and for both haplotypes combined. **c,** Visualization of unphased and phased chromatin contacts in a 14 Mb region on chromosome 7 (chr7: 133-147 Mb). Differential contact count and hotspot regions based on permutation test results are visualised per haplotype. Below, location of H3K27me3 and H3K27ac peaks and ASE genes expressed in H1 hESC. **d,** Imprinted genes and ASE genes overlapping (red) and non-overlapping (black) identified hotspot regions (haplotype 1: red, haplotype 2: yellow). Location of hotspot regions for both haplotypes combined (grey), H3K27me3 (blue gradient) and H3K27ac (green gradient) peak density (counts per bin, 0-99th percentile), and LADs (blue) visualised in vertical tracks. **e,** Percentage of hotspot regions overlapping and non-overlapping H3K27me3 and H3K27ac peaks and LADs. Significance level was determined by performing circular permutations of H3K27me3 and H3K27ac peaks and LADs.

Next, we defined the differential contact count as the number of differential contacts per genomic window (≤10 Mb genomic distance), providing a measure of the local density of haplotype-specific contacts (**Fig. 4a**, **Fig. S6** for all autosomes). Using this metric, we defined hotspot regions as genomic windows exceeding a predefined threshold of differential contact count, thereby further condensing information and highlighting regions with a high enrichment of haplotype-specific contacts (based on Beagrie et al., 2023^20^) (**Fig. 4c**, **Fig. S7a**). We found that ∼15.23% of the genome forms contact hotspots, indicating extensive haplotype specific structural differences in chromatin conformation.

### Haplotype-specific chromatin contacts are enriched for H3K27me3 occupancy in H1 hESC

To begin exploring haplotype-specific differences in chromatin structure, we compared cophased contact matrices across an imprinted gene region, and confirmed haplotype-specific 3D structural differences previously reported using Hi-C^9^ (**Fig. S8**). We then explored in more detail a region on chromosome 7 characterized by a cluster of contact hotspots (chr7:134 - 144 Mb; **Fig. 4c**). In this region, one haplotype shows increased contact intensities at two distinct loci at ∼135-136 Mb and at ∼143.5 Mb which are depleted in the other haplotype. Differential contact hotspot regions were previously shown to be enriched for H3K27me3 occupancy, a repressive histone modification instigated by Polycomb Repressor Complex 2 (PRC2), in F123 mESCs^10^. To investigate whether PRC2 is associated with haplotype-specific contacts in H1 hESC, we acquired publicly available H3K27me3 ChIP-seq collected from the same cell line. In addition to H3K27me3, we also considered H3K27ac, a histone mark associated with active enhancers, and lamina-associated domains^22^ (LADs) as a feature of higher order nuclear organisation. To assess the connection between haplotype-resolved 3D genome organization and gene expression, we also investigated the presence of imprinted and allele-specific expression (ASE) genes. We found that the region ∼135 - 143.5 Mb on chromosome 7 shows prominent haplotype-specific hotspots which are enriched for H3K27me3. Interestingly, the region contains the mono-allelically expressed gene *CYREN* (chr7:135.1 - 135.2 Mb), a gene relevant in cell-cycle-dependent regulation of cell transitions into G or S state.

Based on the observation of individual imprinted and ASE genes overlapping with genomic regions showing differential contacts, we analyzed the general overlap of imprinted and ASE genes with hotspot regions, using two gene lists filtered for ASE and imprinted genes expressed in H1 hESC line as well as two external imprinted gene lists^23,24^. Hotspot regions overlap imprinted and ASE genes across the genome, in some cases in clusters as seen for example on chromosome 15 or chromosome 1 (**Fig. 4d**). Approximately 20-30% of ASE and imprinted genes overlap with hotspot regions (**Fig. S7c**). Repressive Polycomb-mediated facultative heterochromatin marked by H3K27me3 and active enhancer regions marked by H3K27ac show high enrichment for and significant colocalization with hotspot regions (29.3% and 16.8%, respectively), as assessed using circular permutation of peak and domain regions (**Fig. 4e, Fig. S7d,e**). LADs showed a higher but non-significant fraction of overlapping hotspot regions (49.1%) (**Fig. 4e, Fig. S7f**).

### Topological associating domains are highly haplotype-specific in H1 hESC

To investigate differences in higher-order chromatin structure between the two H1 hESC haplotypes, we identified TAD boundaries in the unphased and cophased GAM matrices based on the normalized insulation score^25,26^. A total of 4,783 TAD boundaries were identified genome-wide across unphased and cophased H1 hESC matrices, with similar numbers of TAD boundaries across the three groups (mean = 2,333 TAD boundaries; **Fig. 5a**). While many boundaries are shared among all three matrices (n = 1138), reflecting common structural features, we revealed differences in TAD boundaries between cophased and unphased data, for example in a 5 Mb region with hotspot regions overlapping the imprinted gene *PLAGL1* (**Fig. S9**). A considerable fraction of boundaries is unique to a single haplotype (haplotype 1: 817; haplotype 2: 829), indicating substantial haplotype-specific variation in domain organization in human genomes, as previously reported in F123 mESCs^10^. Additional sets of TAD boundaries were shared between one haplotype and the unphased matrices (haplotype 1: 577; haplotype 2: 587), further supporting structural differences between the two homologous chromosomes at the level of TAD organization.

**Fig. 5:**
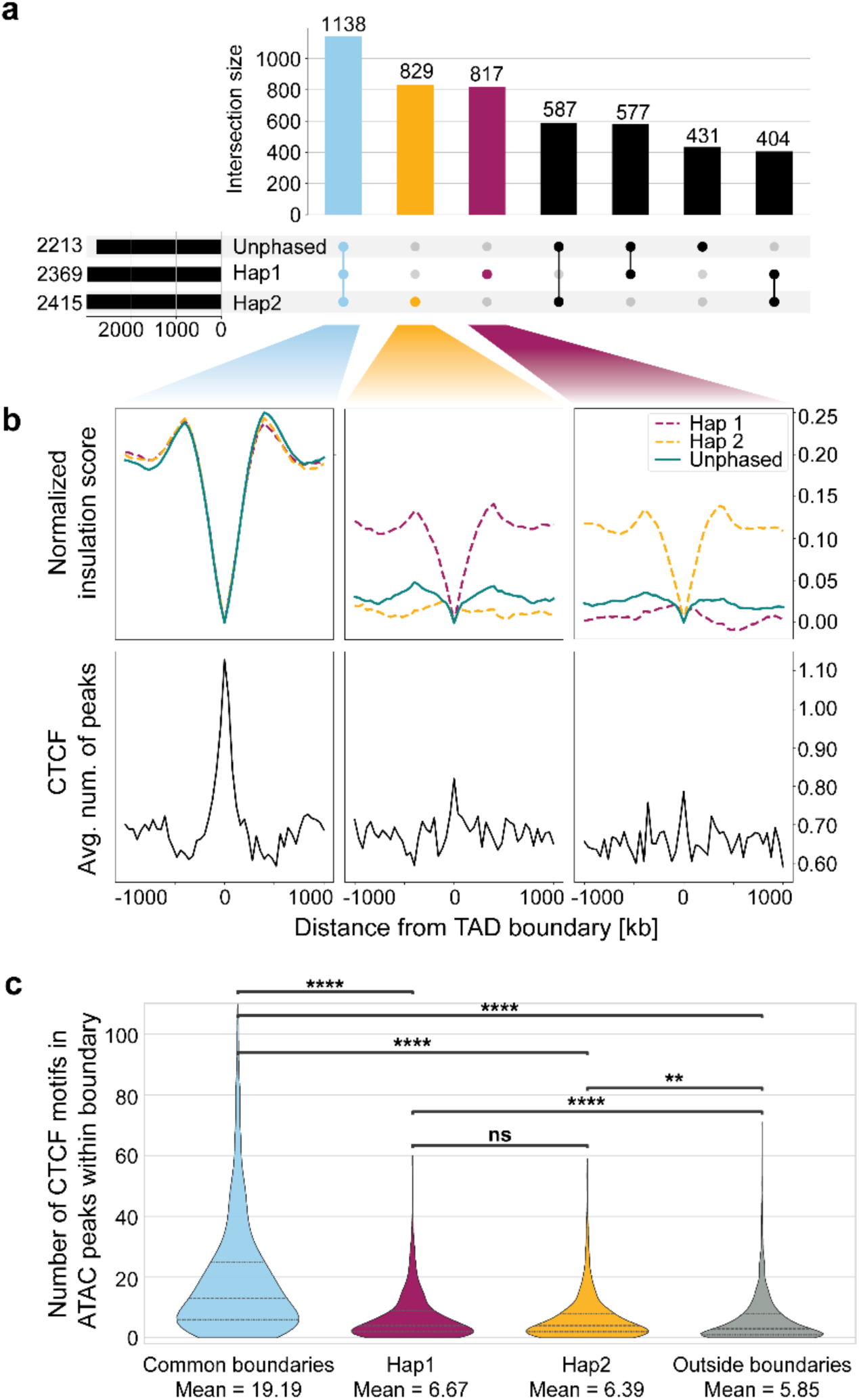
Haplotype-specific TAD boundaries in H1 hESC revealed by CoPhasing. **a,** Intersection of TAD boundaries across unphased and cophased data. **b,** Insulation score average profile for common TAD boundaries (unphased and cophased data), cophased haplotype 1 exclusive TAD boundaries and cophased haplotype 2 exclusive TAD boundaries in the first row. The color of the profiles indicates the normalized insulation score data for unphased data (teal) and cophased data (haplotype 1: red, haplotype 2: yellow). Average profile of CTCF peaks at common TAD boundaries, cophased haplotype 1 exclusive TAD boundaries and cophased haplotype 2 exclusive TAD boundaries in the second row. The profiles are visualized in a window centered at the TAD boundary ±1000 kb. **c,** CTCF binding motifs in ATAC peaks within haplotype exclusive TAD boundaries, common boundaries and outside of boundaries.

At shared TAD boundaries, the average normalized insulation score profiles in unphased and cophased data are highly similar (**Fig. 5b** upper row). Haplotype-unique TAD boundaries show prominent decline in the normalized insulation score profile in each haplotype from which the exclusive TAD boundaries originate, confirming the clear separation of TAD boundaries. Notably, CTCF ChIP-seq peaks are highly enriched at shared TAD boundaries suggesting that CTCF contributes to the formation and stabilization of these boundaries (**Fig. 5b** lower row). In contrast, haplotype-specific TAD boundaries have lower CTCF enrichment levels. Common TAD boundaries contain substantially more CTCF-binding motifs within chromatin-accessible regions than haplotype-specific boundaries, identified using published ATAC-seq data of H1 hESC^27^ (**Fig. 5c**). Nevertheless, haplotype-specific TAD boundaries show significant enrichments for CTCF-binding motifs relative to background levels.

Gene Ontology (GO) enrichment analysis for genes found at the common TAD boundaries showed significant enrichment for genes involved in metabolic and catabolic processes, response to oxidative stress and signalling pathways (**Fig. S10**, **Table S1**), pointing to the general housekeeping genes previously shown to be enriched at TAD boundaries^28^. Genes found at the haplotype-specific TAD boundaries show enrichments for GO terms related to cell adhesion (*RHOA, CDH13, CLDN7*), cell to cell signalling (*LGR5, AXIN2, EGF, LRP5, LRP6*), including cell cycle related genes (*CCND1, CCNE1, CCNY*) and early development factors (*FGF10, TGFB1, PITX2, PAX8*).

In summary, CoPhasing of GAM data reveals haplotype-specific differences in chromatin architecture across the human genome in H1 hESC. Through permutation testing and integration with insulation score, ASE analysis, and chromatin marks, the CoPhasing approach exposes structural differences between homologous chromosomes, advancing our capacity to investigate diploid genome organization in genomes with low SNV density at unprecedented resolution.

## Discussion

CoPhasing of GAM data introduces a conceptual shift in haplotype-resolved chromatin conformation analysis. It leverages the inherent haplotype fidelity of GAM to overcome the limitations of direct phasing approaches that depend exclusively on read-SNV overlaps. CoPhasing markedly improves the phasing efficiency on both the read and the window level, ultimately enabling the construction of high-resolution, haplotype-specific chromatin contact matrices irrespective of SNV density.

To identify differential haplotype-specific chromatin contacts in cophased human GAM data, we developed a permutation test-based algorithm that accounts for spatial dependencies in 3D chromatin contact maps. Alternative approaches, such as distance-normalized contrasts^9^ or z-score subtraction^10^ can be confounded by these dependencies, violating independence assumptions and inflating false positives. Our method mitigates this via post-processing steps including log-transformation, Gaussian filtering, and adaptive thresholding. While further parameter tuning could improve sensitivity, the current implementation provides robust and interpretable results.

Application of this algorithm on the cophased chromatin contact matrices generated from human GAM data revealed extensive haplotype-specific differences in chromatin interactions, genome-wide. While analysis parameters might affect the exact reported fraction, our findings consistently indicate widespread allelic structural variation in the pluripotent H1 hESC.

Our results confirmed the previously reported differential contacts at the imprinted *SNRPN* locus on chromosome 15, a genomic region associated with Prader-Willi syndrome, validating the biological and clinical relevance of the CoPhasing approach on GAM data^9,29^. While human imprinted genes may show variable allelic expression between hESC lines, *SNRPN* and its neighboring imprinted genes *NDN* and *MAGEL2 are* consistently monoallelically expressed across multiple hESC lines^23,24^. Beyond known imprinted loci, CoPhasing revealed extensive, previously undescribed differences in chromatin contacts between haplotypes in a Polycomb-enriched region on chromosome 7, which includes the ASE gene *CYREN*, a cell-cycle regulator of non-homologous end joining^30^.

At the genome-wide level, the hotspot analysis shows that 20% of genes that are allele-specifically expressed in H1 hESC, or known to be imprinted in humans, are located in regions exhibiting abundant differential contacts. To our knowledge, the majority of these genes have not been reported to show haplotype-specific chromatin conformation. Although imprinted genes provide a starting point for investigating haplotype-resolved data, our results suggest that other regulatory mechanisms beyond imprinting are associated with haplotype-specific contacts in hESCs, as previously indicated in Richer et al., 2023^9^. Haplotype-specific chromatin configurations may provide the underlying structural framework for ASE, offering a mechanistic explanation for recent observations of diverse and variable ASE patterns across healthy human tissues^31^. Intriguingly, in contrast to the housekeeping gene enrichment observed at common TAD boundaries, haplotype-specific TAD boundaries are enriched for genes with roles in cell adhesion, cell signalling, and development in the pluripotent H1 hESC, suggesting that allele-specific chromatin architecture may play a role in coordinating the spatiotemporal regulation of gene expression programs during early development and differentiation. Supporting a role for chromatin state in driving haplotype-specific differences, a notable fraction of differential-contact hotspot regions overlaps H3K27me3 peaks, and less prominently, H3K27ac, highlighting the contribution of Polycomb-mediated facultative heterochromatin to haplotype-specific differences, shown previously in mESCs^10^.

Further studies are required to elucidate the interplay of specific epigenetic features, such as asymmetric histone modifications, transcription factor binding or differential DNA methylation, with haplotype-specific chromatin structures presented here. While our current analysis identified overlaps between differential contacts and known imprinted regions, the unknown parental genomes of H1 hESC prevent a deeper exploration of directionality, and limit the distinction between parent-of-origin effects and sequence-driven variation. Additionally, investigating how haplotype-specific contacts vary with genetic variation, across different cell types and developmental stages will be essential to fully understand their role in gene regulation and gene expression variation in the diploid human genome.

While the current application of CoPhasing was restricted to embryonic stem cell lines, which may not reflect the epigenetic landscape of cells in vivo^32^, CoPhasing is applicable to GAM data from other mammalian cell types or tissues. It therefore offers a promising opportunity to investigate haplotype-specific 3D chromatin folding in other contexts, including development, disorder and disease. The integration with emerging single-cell and multimodal approaches will enable direct studies of allele-specific genome organization and its functional impact in patient-derived cells and tissues. Together with GAM’s capacity for haplotype reconstruction^18^, CoPhasing now enables fully haplotype-resolved chromatin architecture mapping, and establishes a foundation for connecting allelic differences in 3D genome organization with gene regulation, epigenomic diversity, and phenotypic variation in health and disease.

## Data and Methods

### Genome Architecture Mapping datasets

Publicly available GAM data from the S129/Jae and Cast hybrid mouse embryonic stem cell line F123^10,33^ was obtained from GEO (GSE25471, Table S2). Publicly available GAM data from the human embryonic stem cell line H1^3,34^ was obtained from the 4DN data portal^35^ (4DNESVAMUDHA, 4DNES44S71I5), also available from GEO (GSE324313).

### Haplotypes

Phased F123 mESC SNVs were obtained from Irastorza-Azcarate et al., 2025^10^. Phased H1 hESC SNVs were available from Dixon et al., 2015^8^ based on hg18. H1 SNVs were lifted over to GRCh38 using UCSC liftover tool^36^.

### Direct phasing of GAM data

Phased GAM data from F123 mESCs was available from Irastorza-Azcarate et al., 2025^10^ (GSE254717). Briefly, GAM sequencing data is mapped to the N-masked genome, where all SNVs positions are replaced by N, using default parameters of bowtie2^37^ (version 2.3.4.3). The reads mapped to the N-masked genome are checked for the presence or absence of SNVs, and sorted to the haplotype-specific bam-files with SNPsplit package. Next, the genome is split into equal-sized windows and the optimal threshold of nucleotide coverage for calling positive windows is calculated using the coverage per bin from all reads. Windows are phased to a haplotype when the number of nucleotides covered by the reads containing haplotype-specific SNVs is higher or equal to the optimal threshold.

### CoPhasing pipeline

The CoPhasing pipeline that implements the developed CoPhasing strategy is available as a nextflow pipeline in the GitHub repository (https://github.com/ICCB-Cologne/cophasing). The pipeline (**Fig. S1**) requires three input file types: i) BAM files of a GAM experiment, ii) FASTA file with the appropriate reference genome, and iii) VCF file with phased SNV information. In addition to the input files, parameters for the pipeline can be defined in a config file. Parameters for the pipeline are the name of the output directory (out), name of the output directory for intermediate files (debug_out), different resolution values to be used (bins = [40000, 50000, 100000]), CoPhasing distance threshold (cutoff), a minimum required read depth per SNV to be considered (min_depth), and a required minimum ratio of observed bases (min_ratio). For all parameters except for the debug_out parameter a default value is set in the pipeline. If the debug_out parameter is defined, intermediate output files will be saved in this directory to enable quality control and process monitoring of each step in the pipeline. The main output files, coverage tables and segregation tables, are saved in the defined output directory. The coverage table is a matrix of genomic windows using set resolution values (chromosome, start, stop) by sample indicating read counts of the sample in the region. The segregation table is a binary matrix with the same structure indicating the presence of a region in samples. In the output directory both the coverage and the segregation tables (.tsv) for each defined resolution value for both haplotypes (“hap1”, “hap2”) and the unphased results (“both”) are stored for further analysis.

### Quality of genome sampling in non-phased and cophased GAM data

The suitable working resolution of the pairwise GAM contact matrices were determined from the frequency of genome sampling quality, as described previously^10^. Briefly, the distribution of raw co-segregation events for all intra-chromosomal pairs of genomic windows was compared to the standard Poisson distribution, with the same mean and standard deviation, using a non-parametric Kendall rank correlation coefficient. Kendall’s τ correlation coefficient ≥ 0.95 was considered as the indication of good quality of genome sampling at a specified resolution and genomic distance.

### Identification of differently sampled regions in GAM segregation data

To identify regions with robust sampling in the GAM process, we computed the window detection frequency (WDF) as the ratio of NPs in which a genomic window was detected over the total number of NPs in the segregation table.

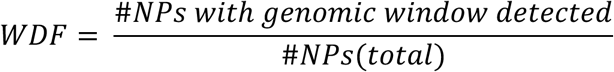

Genomic windows were classified as undersampled or oversampled by defining upper and lower bounds as the mean WDF per chromosome ± a constant factor (*f*) multiplied with the standard deviation:

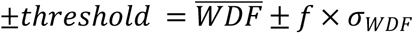

Genomic windows with a WDF more extreme than the upper or lower threshold or with no detection events (WDF=0) were excluded from further analysis. For the segregation tables of two haplotypes, the regions were excluded in both haplotypes, if excluded in either one of them.

### Computation of normalized pairwise GAM contact matrices using NPMI

Pairwise GAM contact matrices were computed using normalized pointwise mutual information (NPMI) for all pairs of genomic windows, as previously described^26^. NPMI describes how the joint probability of a pair of genomic windows being found in the same NP deviates from the expectation based on their individual probabilities across all NPs.

### Parameter optimization for generation of high-resolution cophased contact matrices

The high SNV density of F123 mESC was leveraged to benchmark the CoPhasing approach and to determine the optimal resolution and distance thresholds that maximize information gain while maintaining high accuracy. To simulate human SNV density, the F123 SNV set with an average SNV distance of 132 bp was downsampled to 2 kb by random subsampling. The mm10 genome size in bp was divided by 2000 bp to calculate the target number of SNVs for the desired average genomic distance of 2kb. From the full F123 set of 18,150,228 SNVs, 6.8% were retained for the downsampled set of 1,231,373 SNVs using the R function sample() without replacement. (**Fig. S2**). Comparison of 10 different iterations of subsampled SNV sets yielded highly comparable results, only one was reported here. We assessed the impact of increasing CoPhasing thresholds (0 bp as direct phasing baseline, 1 kb, 50 kb, 100 kb, 200 kb, 1 Mb, 10 Mb, and full chromosome length) on the efficiency and accuracy of haplotype-assignment on the read, window, and matrix level by comparing to respective results from the same GAM dataset directly phased on the full F123 SNV set.

### Insulation score calculation and TAD border calling

Insulation score (IS) tracks were calculated from NPMI GAM contact matrices with 40 kb resolution using the insulation square method as previously described^25,26^. A sliding window (insulation square) of a given size is shifted along the diagonal of the matrix to calculate the mean contact frequency for each window. The results of the different insulation square sizes (80 kb, 160 kb, 240 kb, 320 kb, 400 kb, 480 kb, 560 kb, 640 kb, 720 kb, and 800 kb) were visualized stacked with 80 kb on the bottom row and 800 kb on the top. TAD borders were called using a 400 kb insulation square size and based on local minima of the normalized insulation score with one genomic bin added on each side.

### Permutation test to identify differential haplotype-specific contacts

To identify significant differences in 3D chromatin conformation between homologous chromosomes a permutation test was applied to the differences in contact intensities between the two cophased datasets (**Fig. S5**). To this end, chromatin contact matrices were calculated displaying contact frequencies for the two haplotypes separately based on each cophased segregation table. Differences between the two haplotypes (ΔCF) were calculated by subtracting contact frequencies for Hap2 from Hap1. To generate the null distribution the cophased segregation tables of both haplotypes were concatenated into a joint segregation table and randomly split without replacement into two permuted phased segregation tables, Hap1’ and Hap2’. ΔCF’ scores were then computed as described for the observed data. These two steps were repeated N = 2000 times to generate a null distribution, against which the observed ΔCF values were evaluated using an empirical p-value:

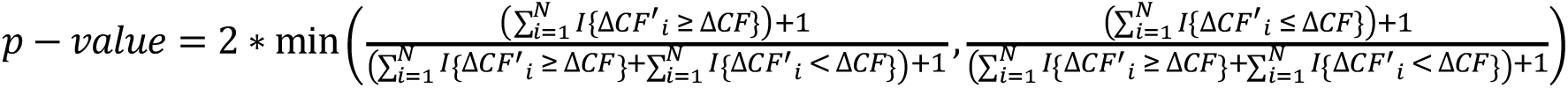

Owing to the non-independence of tests due to spatial effects, p-values were log-transformed (-log_10_) and underwent spatial smoothing and thresholding. To smooth a matrix containing missing values, a normalized Gaussian convolution was applied. First, a binary mask was generated with missing values set to 0 and valid entries set to 1. Second, the matrix with missing values replaced by zeros and the mask were smoothed with a Gaussian filter with a determined kernel size (here kernel size = 1). Third, the smoothed matrix was normalized by the convolved mask to correct for missing values and ensure that matrix entries get averaged over valid neighboring matrix entries, preventing NaN propagation. Fourth, in the smoothing procedure, the result was symmetrized by averaging with its transpose and previously missing present NaN were reintroduced. Lastly, a threshold for each chromosome was applied to filter and identify regions in the data with haplotype-specific contacts. The respective thresholds were determined by plotting a range of thresholds (0.05 steps) against the corresponding ratio of elements above the thresholds and identifying the knee point of the curve for each chromosome. Directionality was added to the results based on the information if more permutations show higher or lower ΔCF’ values compared to the original ΔCF at that position. Permutation test results which show stronger contacts on Hap2 were assigned negative scores for matrix visualization of changes between both haplotypes.

### Detection of differential contacts, haplotype-specific contact hotspots and extended genomic regions with increased allelic contact differences

Differential contacts were determined based on the permutation test results. For each genomic bin, the number of differential contacts was then quantified by counting differential interactions within 10 Mb to upstream and downstream regions of the matrix, separately for each haplotype ^20^. Hotspot regions were defined as bins exceeding a threshold of 28 counts of differential contacts, selected based on visual inspection of hotspot signal tracks. To identify extended genomic regions of increased allelic differences at contact level a sliding window approach was applied. The number of differential contacts were summed up in a 10 Mb section and the window was advanced in steps of 1 Mb, resulting in overlapping windows across the genome.

### ASE analysis

H1 expression data was acquired from ENCODE (ENCSR895ZTB). Allele-specific expression of H1 was determined through an analysis pipeline building on GATK ASEReadCounter, available at https://github.com/icgc-argo-workflows/allele-specific-expression.

### CTCF motif enrichments on TAD boundaries

Chromatin accessibility of H1 hESC was determined by ATAC-seq peaks acquired from the 4DN Data Portal^27^ (4DNFIPGM38K4). Chromatin-accessible regions within unphased and haplotype-specific TAD boundaries (120 kb windows) were identified by intersecting boundary coordinates with ATAC-seq peaks using bedtools intersect. CTCF binding motifs located within these accessible regions were then identified using the motif-analysis module of the Regulatory Genomics Toolbox^38^. Motif scanning was performed using the CTCF position weight matrix CTCF_HUMAN.H11MO.0.A from the HOCOMOCO v13 database. The number of motif occurrences within accessible regions was quantified separately for unphased and haplotype-specific TAD boundaries. Background genomic regions were generated using bedtools shuffle (-chrom --noOverlapping), using the union of all detected TAD coordinates from both unphased and cophased datasets as input.

### Gene Ontology (GO) enrichment

GO enrichment analysis of genes was performed using Web Gestalt (https://www.webgestalt.org/). All genes were used as the background universe. Overrepresentation analysis was performed selecting Gene Ontology as a Functional database in the website. The overlap was performed using the TSS coordinates of genes within a 120 kb TAD boundary region centered at the bin with the lowest insulation score.

### Visualization of integrated epigenetic tracks with cophased matrices

For the visualization of chromatin contact matrices for unphased and phased GAM data the limits were set to zero for the minimum value and to the maximum 99th percentile of the three chromatin contact matrices (Unphased, Haplotype 1, Haplotype 2) for maximum value. The color scale ranges from blue for less frequent contacts to red for more frequent contacts between the regions along the linear genome. Filtered regions were identified based on window detection frequency and displayed in light grey (Methods: Identification of differently sampled regions in GAM segregation data). To identify unmappable regions visualized in GRCh38 gaps track, genome mappability for GRCh38 human genome assembly was computed using GEM-Tools suite with a read length of 75 nucleotides, matching the read length of the sequencing data^39^. Mean mappability scores were computed for each genomic bin using bigWigAverageOverBed. Genomic bins with a mean mappability ≤ 0.1 in the human genome were defined as GRCh38 gaps.

The PyGenomeTracks tool was used to visualize CoPhasing results in chromatin contact matrices together with additional information like metrics to evaluate differential contacts, insulation scores, GRCh38 gaps, hotspot regions and genes located in the visualized region.

## Experimental Methods

### H1 hESC culture

H1 human embryonic stem cells (H1 hESCs, WiCell, WA01) were maintained under feeder-free conditions on tissue culture-treated plates (Corning, Cat. #353046 or equivalent) coated with hESC-qualified Matrigel (Corning, Cat. #354277) and cultured in mTeSR1 medium (STEMCELL Technologies, Cat. #85850). Cultures were maintained in a humidified incubator at 37°C with 5% CO2, with 10 mL mTeSR1 per 10cm dish and daily medium replacement. Cells were monitored microscopically each day and passaged approximately every 4-5 days using the non-enzymatic dissociation reagent ReLeSR (STEMCELL Technologies, Cat. #05872) to preserve colony growth as small aggregates rather than single cells. Briefly, cultures were washed with PBS without Ca2+ and Mg2+, exposed to ReLeSR for 5-7 min at 37°C, and detached by gentle agitation before replating onto matrigel-coated plates in mTeSR1 media with ROCK inhibitor Y-27632 (STEMCELL Technologies, Cat. #72304) at an appropriate split ratio of 1:10. Regular feeding was conducted with mTeSR1 without ROCK inhibitor.

### ChIP-seq

Approximately 4 million cells were used for each histone ChIP-seq experiment. Cells were cross-linked with 1% formaldehyde for 10 minutes, followed by quenching with 125mM glycine for 5 minutes and washed for twice with PBS. The cell pellet was lysed in cell lysis buffer (20mM Tris-HCl, pH8, 85mM KCl, 0.5% NP-40) supplemented with 1x protease inhibitor cocktail (Roche, 11836170001) on ice for 20 minutes then pelleted at 2500g for 5 minutes. The pellet was resuspended in sonication buffer (10mM Tris pH7.5, 1% NP-40, 0.5% sodium deoxycholate and 0.1% SDS) with 1x protease inhibitor cocktail and incubated for 10 minutes at 4oC. In order to achieve 200-700nt DNA fragmentation range, nuclei were sonicated (Bronson sonifier model 250 with the following conditions: Amplitude = 10%, (pulse ON/OFF = 0.7sec/1.3sec) for 8-12 min. Chromatin was pre-cleared and incubated with Dynabeads Protein A (Invitrogen, 10002D) for 1 hour. Samples were incubated with 2l antibody overnight at 4C (anti-H3K27me3: Active Motif cat# 39155, batch 31814017; anti-H3K27ac: Diagenode, cat#C15410196, batch A1723-0041D). On the following day, 40l of Dynabeads Protein A were added to the chromatin and antibody mix for 2-3 hours and then the captured immuno-complexes were washed as follows: 3x low-salt buffer, 3x High-salt buffer, 2x LiCl salt buffer and 2x TE. The washed immuno-complexes were eluted in 50l ChIP-DNA elution buffer (10mM TrisHCl pH8, 100mM NaCl, 20mM EDTA and 1% SDS) for 15 minutes each, reverse cross-linked overnight at 65C. DNA was purified using AMPure XP beads, washed three times with 70% ethanol, dried, and eluted in 30 µl elution buffer containing RNase A (Thermo Fischer Scientific, EN0531). The Illumina library construction steps were carried out with 5-10ng of purified ChIP’ed DNA. During the library construction, PCR purification was performed after every step using Qiagen kit (QIAquick PCR purification 28104 or QIAquick gel extraction 28706). The library reaction steps with ChIP’ed DNA as follows: 1) End-repair - T4 PNK (NEB, M0201L), Klenow fragment (NEB, M0210S), T4 DNA Polymerase (NEB, M0203L) and dNTP mix (Thermo Fischer Scientific, 18427013) incubated at RT for 45min, 2) 3’ end A-base addition - 3’ -> 5’ exo- Klenow fragment (NEB, M0212L) and dATP (NEB, N0440S) incubated at 37C for 30min, 3) Adapter ligation - T4 DNA ligase (NEB, M0202M) and illumina adapters incubated at 16C overnight and 4) PCR amplification - 2x PfU Ultra Hotstart master mix (Agilent, 600850-51) for 14 cycles. The amplified libraries were size selected for 200-450nt on 2% Agarose E-gel (Thermo Fischer Scientific, G402002). Paired-end sequencing was performed on Illumina NextSeq 500.

## Data availability

Raw FASTQ sequencing files for all samples from the H1 hESC GAM dataset, together with normalized cophased chromatin contact maps and normalized direct phasing chromatin contact maps have been submitted to the GEO repository under accession number GSE324313. H1 differential contacts as well as CoPhasing results for F123 data with the full set of SNVs are available from Zenodo repository https://doi.org/10.5281/zenodo.19740137. H3K27me3 and H3K27ac ChIP-seq data of H1 is available on the 4DN portal under accessions 4DNFIGKR9C4F, 4DNFI9U2V6V4 and 4DNFIKG1KI8B, 4DNFI73NTNGZ.

## Resources

| Data | Source |
| --- | --- |
| F123 GAM dataset | 4DN data portal<sup>35</sup>: 4DNESRQDNG61,<br>Generated by Ana Pombo lab<br>Irastorza-Azcarate et al., 2025 <sup>10</sup> |
| H1 GAM dataset | 4DN data portal<sup>35</sup>: 4DNESVAMUDHA,<br>Generated by Ana Pombo lab<br>GEO: GSE324313<br>Dekker et al., 2026 <sup>3</sup> |
| F123 haplotype | Irastorza-Azcarate et al., 2025 <sup>10</sup> |
| H1 haplotype | Dixon et al., 2015 <sup>8</sup> |
| H1 Gene expression | ENCODE<sup>40</sup>: ENCSR895ZTB<br>GEO: GSE175297 |
| Imprinted genes | https://www.geneimprint.com/site/genes-by-species.Homo+sapiens.imprinted-All<br>Frost et al., 2011 <sup>23</sup>,<br>Kim et al., 2007 <sup>24</sup> |
| CTCF ChIP-seq | CistromeDB<sup>41</sup>: 58481<br>GEO: GSM1705254<br>Ji et al., 2016 <sup>42</sup> |
| H3K27me3 | 4DN data portal<sup>35</sup>: 4DNFIGKR9C4F, 4DNFI9U2V6V4,<br>Generated by Rene Maehr lab |
| H3K27ac | 4DN data portal<sup>35</sup>: 4DNFIKG1KI8B, 4DNFI73NTNGZ,<br>Generated by Rene Maehr lab |
| LADs | 4DN data portal<sup>35</sup>: 4DNFIJFD4TWE,<br>Generated by Bas van Steensel lab,<br>Kumar et al., 2024 <sup>22</sup> |
| ATACseq | 4DN data portal<sup>35</sup>: 4DNFIPGM38K4,<br>Generated by Rene Maehr lab<br>Akgol Oksuz B et al, 2021 <sup>27</sup> |

## Competing Interests Statement

A.P. holds a patent on ‘Genome Architecture Mapping’: Pombo, A., Edwards, P.A.W., Nicodemi, M., Scialdone, A. and Beagrie, R.A. Patent PCT/EP2015/079413 (2015).

## Acknowledgements

The authors acknowledge support from the DFG Priority Program SPP2202 ‘Spatial Genome Architecture in Development and Disease’, SPP2202 (Project number 422841138), A.P. acknowledges support from the Helmholtz Association, the National Institutes of Health Common Fund 4D Nucleome Program grants U54DK107977 and 1UM1HG011585-03, funded by the Deutsche Forschungsgemeinschaft (DFG, German Research Foundation) – 545980026 (Leibniz Prize), and the Johns Hopkins Bloomberg Distinguished Professorship funds.

R.F.S. is a Professor at the Cancer Research Center Cologne Essen (CCCE) funded by the Ministry of Culture and Science of the State of North Rhine-Westphalia. This work was partially funded by the German Ministry for Education and Research as BIFOLD - Berlin Institute for the Foundations of Learning and Data (ref. 01IS18025A and ref 01IS18037A).

We furthermore thank the ITCC (IT Center University of Cologne) for providing compute resources on the DFG-funded HPC (High Performance Computing) system RAMSES (Research Accelerator for Modeling and Simulation with Enhanced Security) as well as support (DFG funding number: INST 216/512-1 FUGG).

R.M. acknowledges support from the National Institutes of Health [DP3 DK111898 and UM1 HG011536]. The authors thank J. Huey for ChIP-seq data curation.

## Author contributions

J.M. and R.F.S. conceptualized the CoPhasing method. A.S., J.M., A.K. participated in building the CoPhasing pipeline. J.M. and A.K. performed benchmarking and investigated parameter optimization. C.R. and R.F.S. conceptualized the permutation testing to identify differential contacts. C.R. and C.J.T. performed differential contact analysis and hotspot detection. K.M.P. conducted H1 hESC cell culture and cell harvest in preparation for ChIP-seq performed by H.M. under supervision of R.M.. C.R., A.K., J.M. analyzed matrices and curated, integrated, and analyzed genetic and epigenetic data. A.K. and C.R. prepared data submission. J.M., A.K., C.R., and C.J.T. wrote the initial draft with edits from A.P. and R.F.S.. A.P. and R.F.S. supervised the project.

## Supplementary Figures

**Fig S1:**
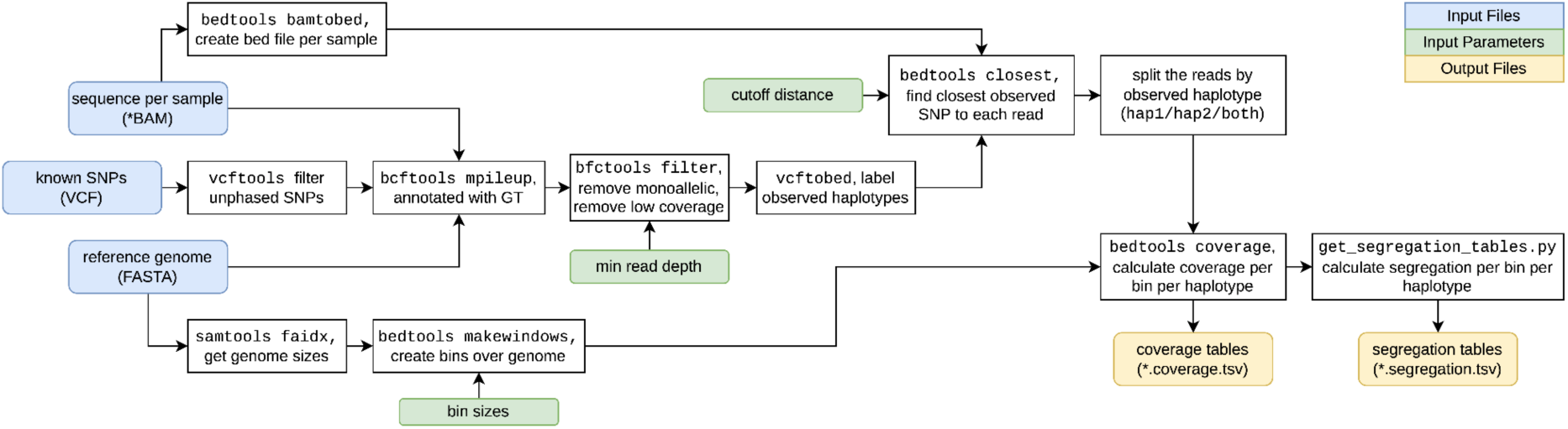
Overview of the CoPhasing pipeline. The CoPhasing pipeline takes mapped reads for each NP of a GAM dataset, phased SNVs of the sample, and a reference genome as input. Using *bcftools pileup*, all observed alleles at SNV positions are detected per NP and further filtered with *bcftools filter* to remove monoallelic sites and low-coverage positions (controlled by the **min read depth** parameter). For each NP, mapped reads and SNVs with observed haplotype information are converted to bed format using *bedtools bamtobed* and *bcftools*, respectively, and *bedtools closest* is used to determine for each read its nearest observed SNV and their distance, which is used to filter by the CoPhasing distance threshold (controlled by **cutoff distance** parameter). Depending on the observed haplotype of their nearest SNV, reads are sorted into phased .bam files. The genome is binned into contiguous, non-overlapping genomic windows using *bedtools makewindows* (resolution controlled by **bin size** parameter), and the coverage of cophased reads in each genomic window is determined using *bedtools coverage*. The coverage information is stored in coverage tables, and used to calculate segregation tables.

**Fig. S2:**
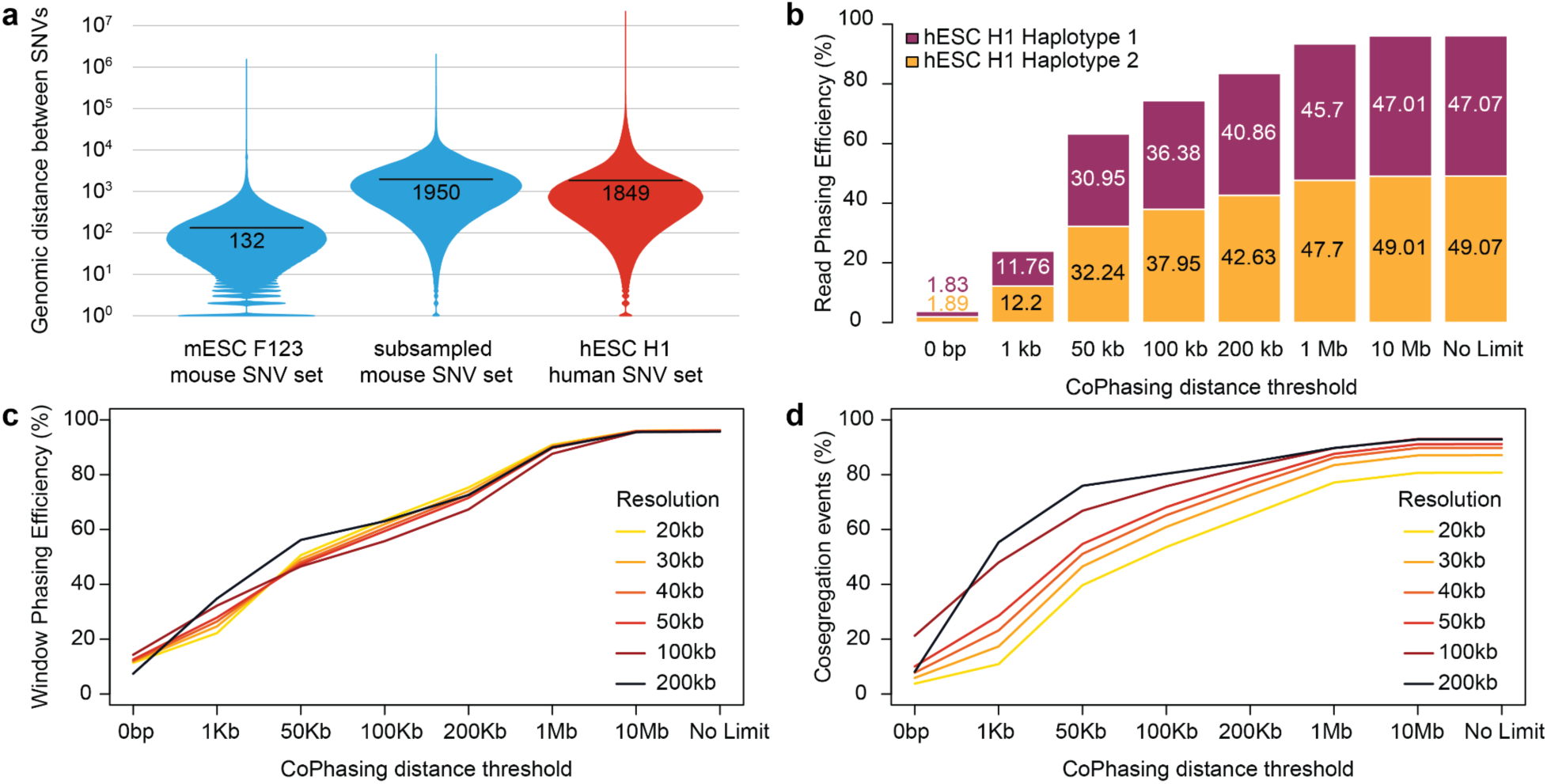
Increased Phasing Efficiency in H1 hESC GAM data. **a,** The dense SNV set of the hybrid F123 mESC line was downsampled from median 132 to ca 2000 SNPs/kb to simulate the SNV density of a human genome, here H1 hESC line. **b,** Applying the CoPhasing approach to the H1 hESC GAM dataset with increased cophasing distance increases read phasing efficiency from below 4% to over 96% of all mapped reads plateauing around 10 Mb distance. **c,** Increased read phasing in the H1 GAM dataset improved phasing of genomic windows with equivalent efficiency at 20 - 200 kb, reaching 100% phasing of all positive windows present in unphased H1 GAM data. **d,** Efficient phasing of genomic windows results in excellent detection of co-segregated pairs of windows, to a level similar to unphased H1 GAM data.

**Fig. S3:**
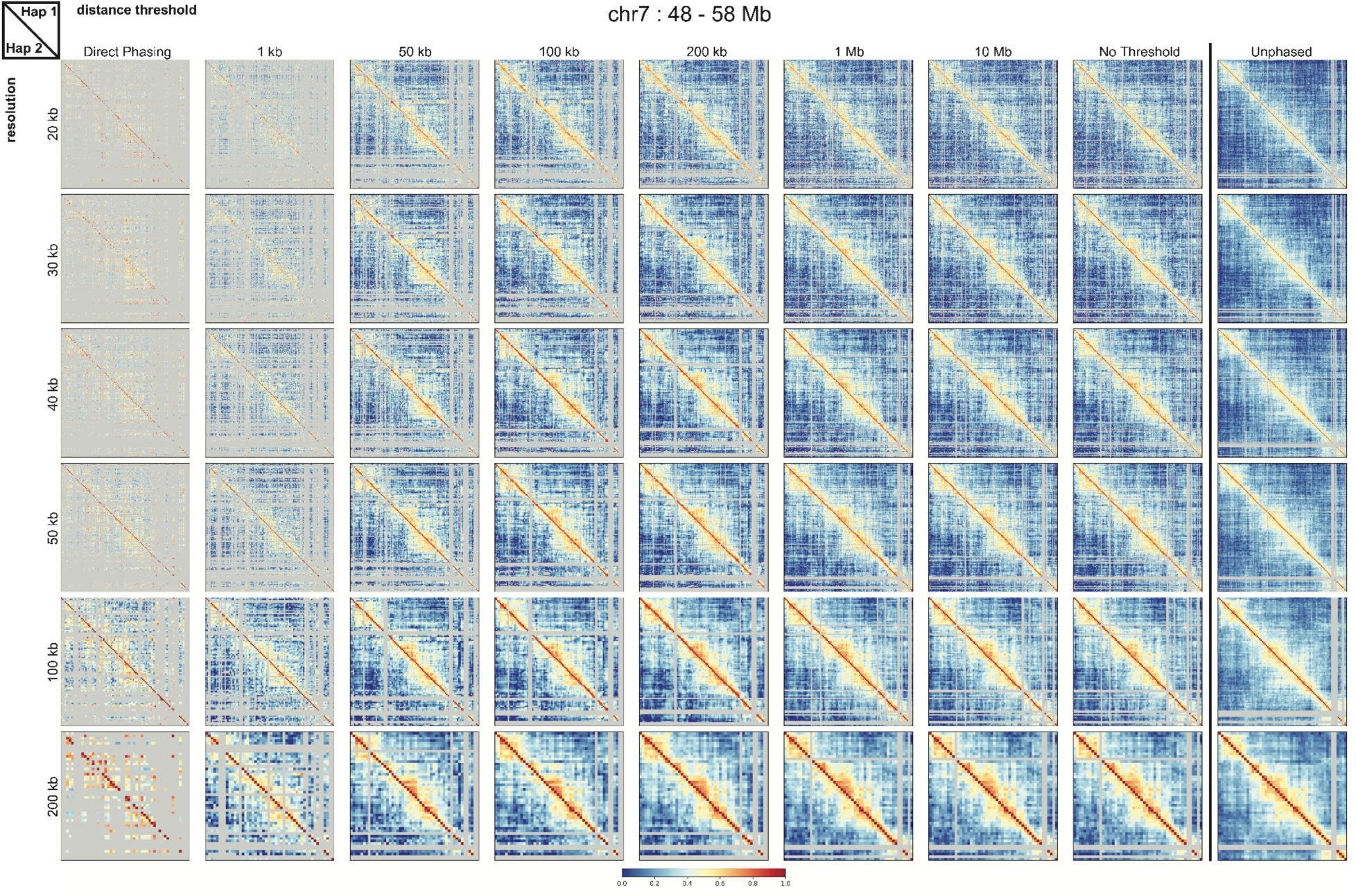
Increased CoPhasing distance thresholds generate well-saturated contact matrices with high resolution from H1 hESC GAM data. Cophased contact matrices were generated for an exemplary 10 Mb region on chromosome 7 (48 - 58 Mb) at increasing resolutions (rows) and CoPhasing distance thresholds (columns). Direct phasing results are shown for comparison in the leftmost column, and unphased matrices are displayed in the rightmost column for each resolution. The upper right and lower left triangles represent haplotype 1 and haplotype 2, respectively, whereas unphased matrices are symmetrical. Contact frequencies are represented by normalized pointwise mutual information (NPMI), with a unified color scale ranging from blue (low contact frequency, 0.0) to red (high contact frequency, 1.0). Grey rows and columns mark under- or over-sampled regions excluded during curation based on window detection frequency (WDF).

**Fig. S4:**
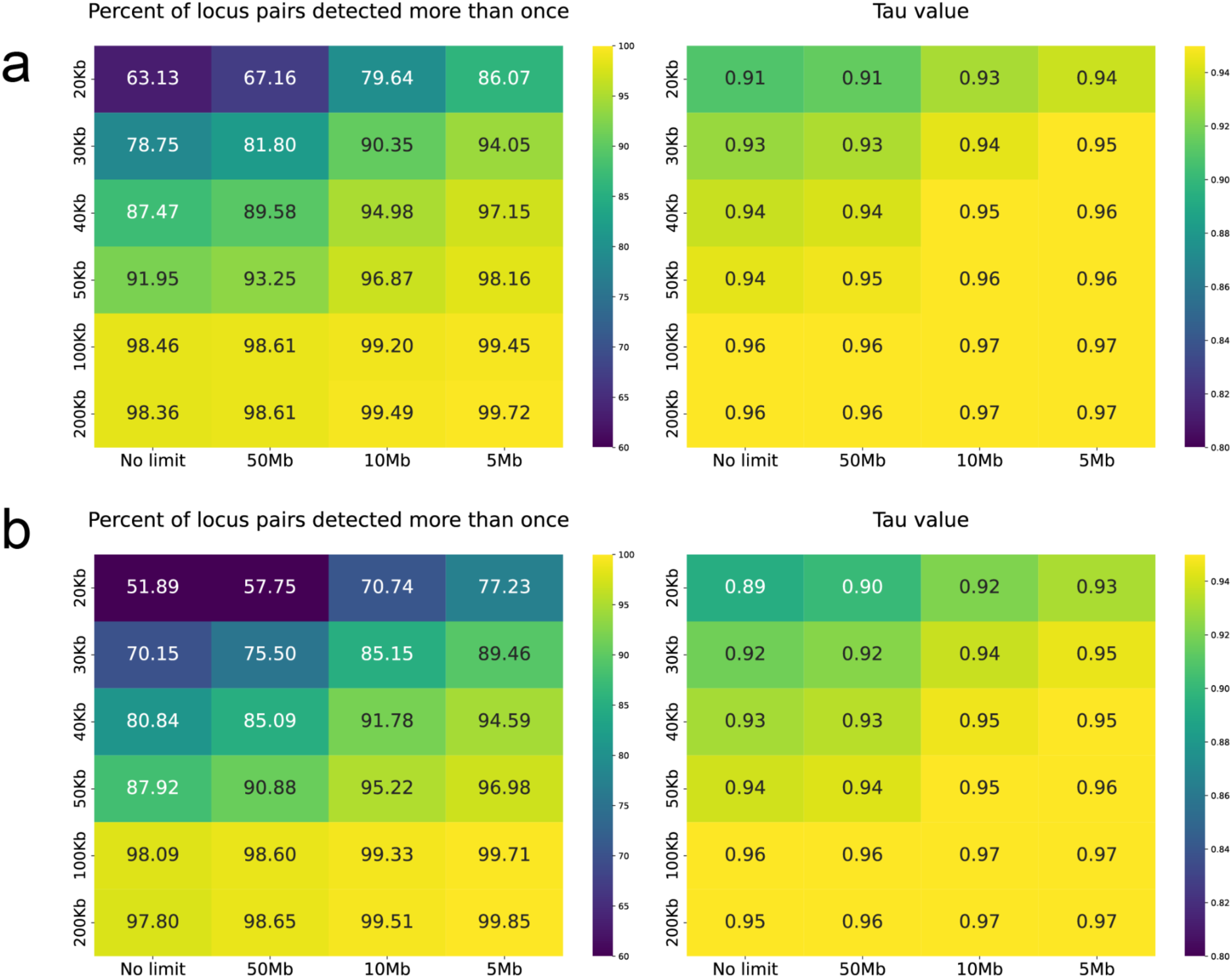
Percentage of locus pairs detected more than once and Kendall’s τ coefficient values for cophased H1 hESC at different resolutions and distances. **a,** Haplotype1 and **b,** Haplotype2. These metrics were used to decide on optimal resolutions and genomic distance for the chromatin contact matrices.

**Fig. S5:**
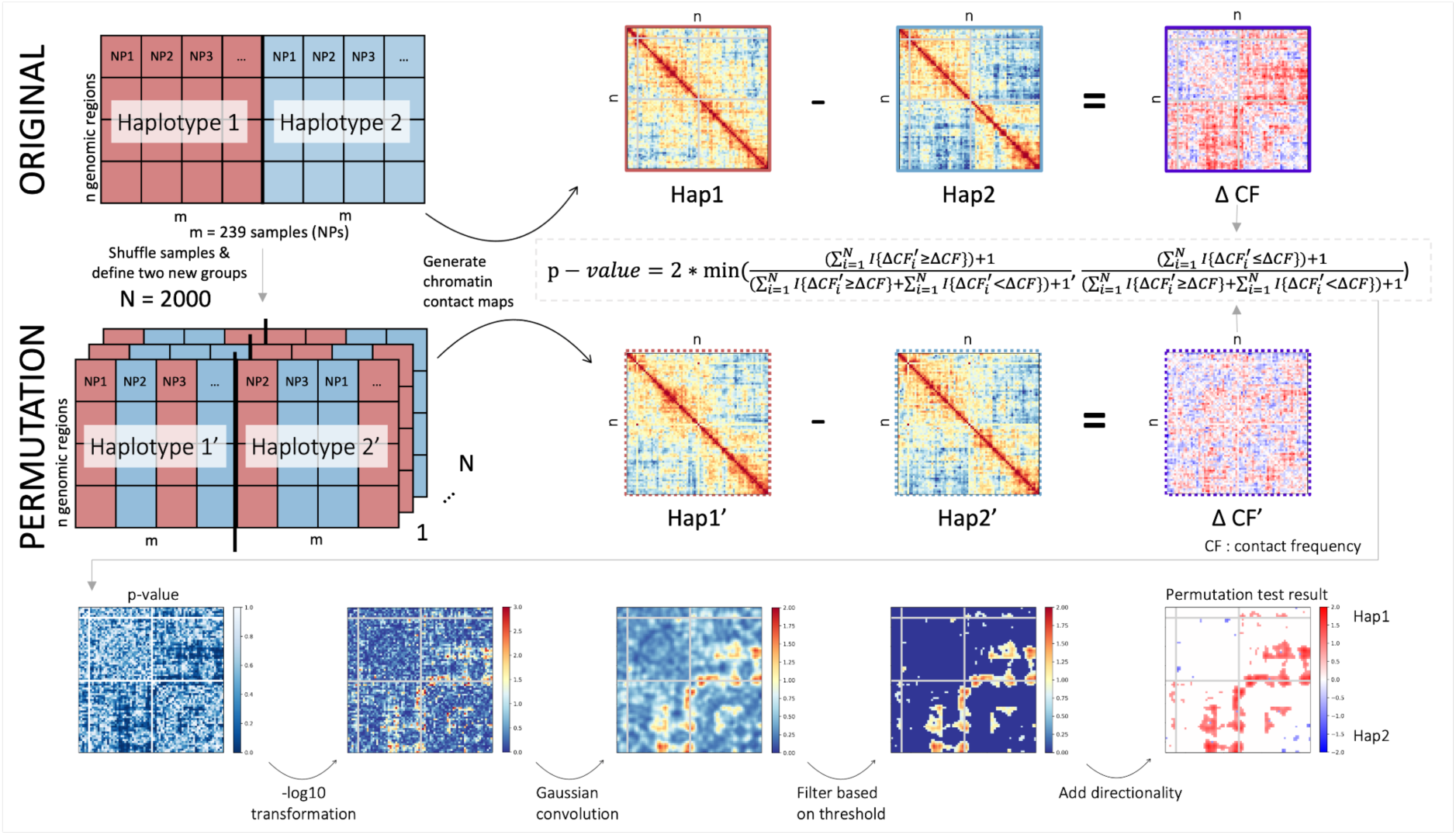
Overview of permutation test workflow for identifying differential contacts. Permutation test workflow to detect significant differential contacts in haplotype-specific chromatin contact maps from GAM data. Differential contact frequencies between both haplotypes ΔCF = Hap1 - Hap2 are calculated from the original dataset obtained from CoPhasing. For 2000 permutations segregation tables for both haplotypes were pooled and randomly split into two permuted haplotypes, yielding ΔCF’ = Hap1’ - Hap2’. The resulting permuted ΔCF’ values are compared to the observed ΔCF and an empirical p-value is calculated for each window. Since the contact frequencies of neighboring windows are often correlated and thus the tests are not independent, a smoothing and thresholding approach is performed to detect regions of differential contacts. Firstly, the p-value matrix is transformed (-log_10_) and smoothed using a gaussian filter with a kernel size of 1. Lastly, the resulting contact values are discretized using an estimated threshold and the directionality of the contact difference is indicated by the sign function.

**Fig. S6:**
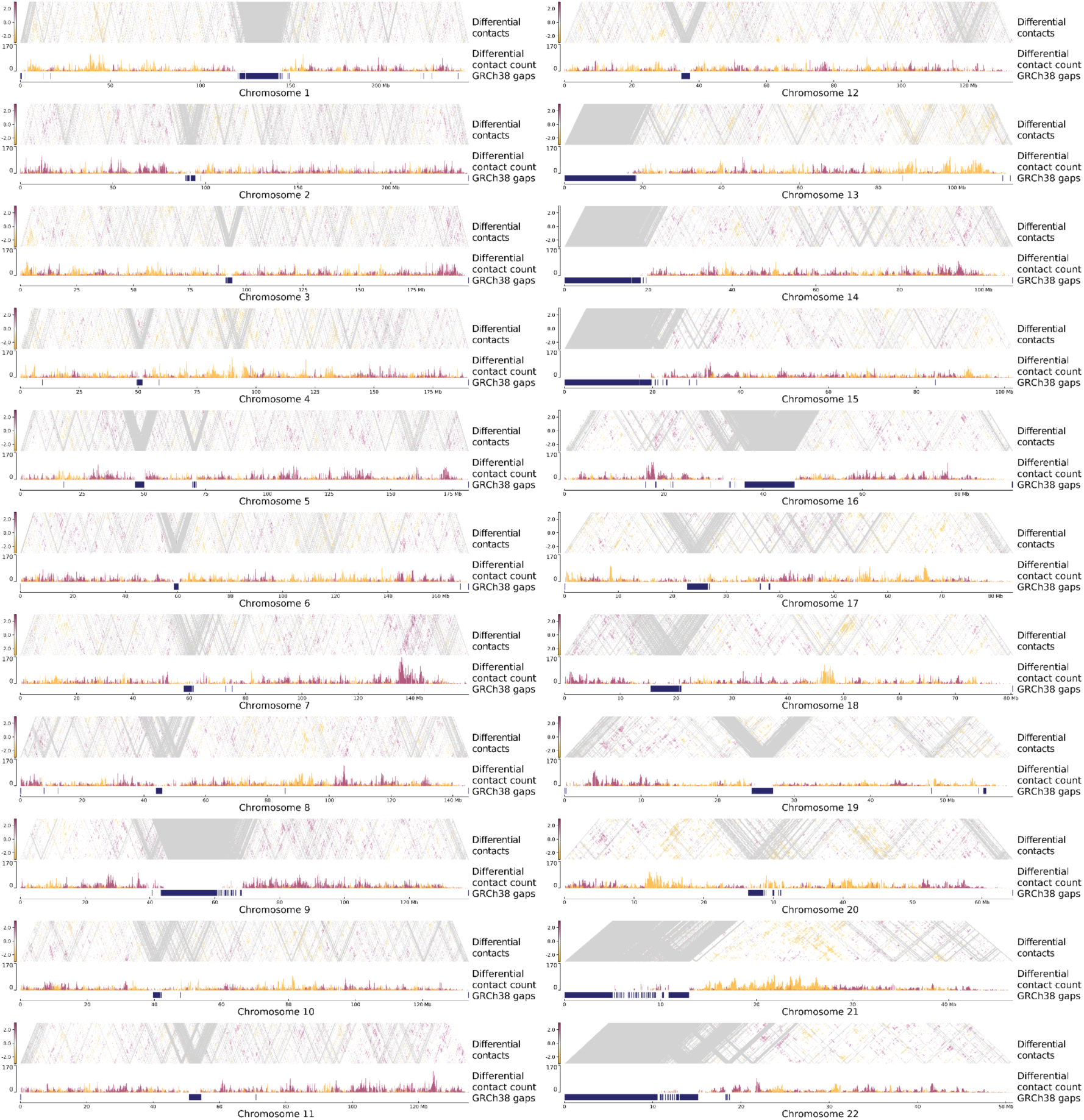
Differential contacts in the H1 hESC genome per chromosome. Three tracks are shown for each chromosome: i) Differential contacts between genomic regions as identified by the permutation test and visualized in a contact matrix. Color scale indicates directionality and intensity of contact (red: stronger contact on Hap1, white: no difference detected, yellow: stronger contact on Hap2). ii) Differential contact count per genomic window within 10 Mb distance (red: contact on Hap1, yellow: contact on Hap2). iii) GRCh38 gaps: Regions with low mappability in the human genome (mappability ≤ 0.1).

**Fig. S7:**
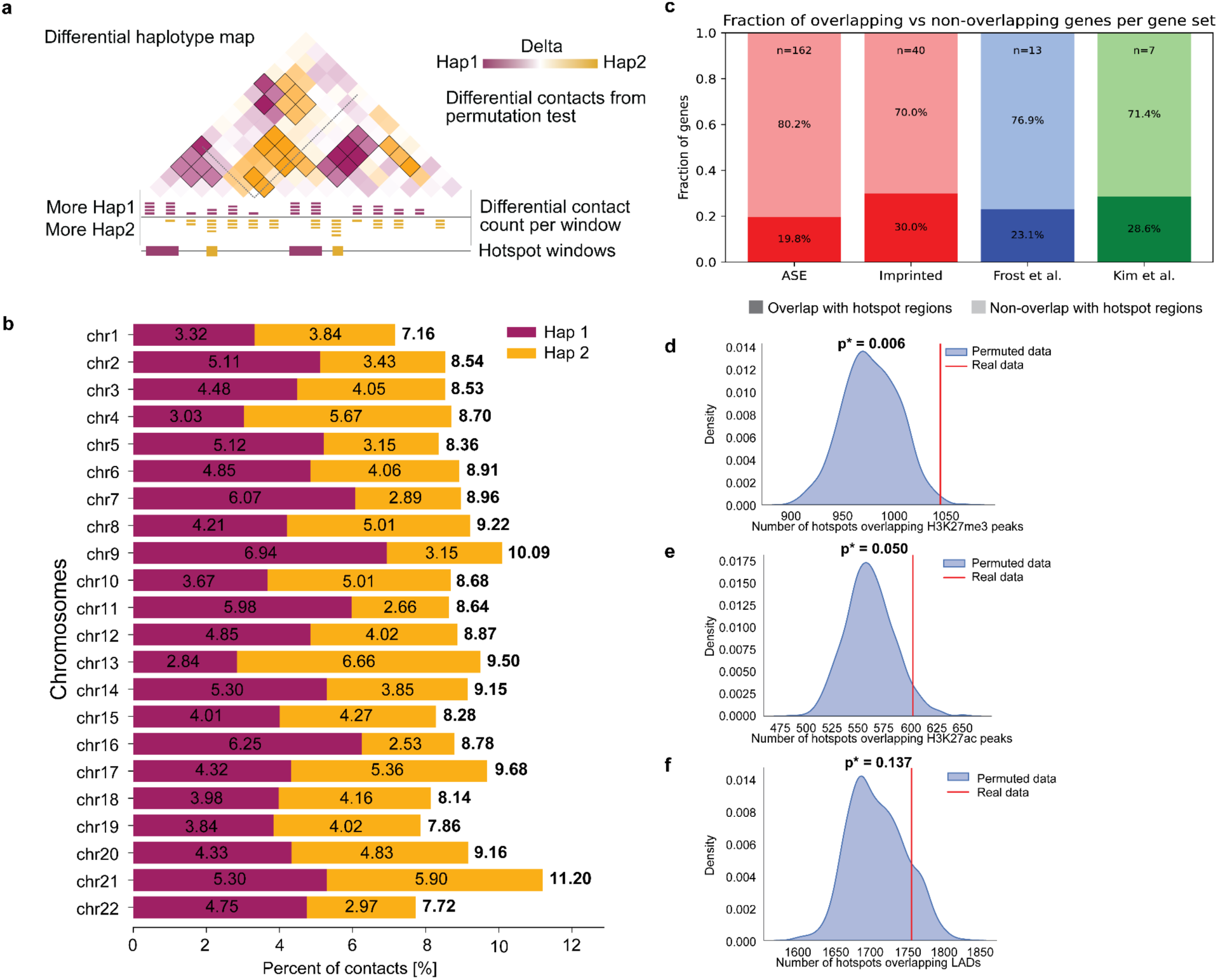
Hotspot analysis based on differential contacts identified by the permutation test reveals regions with strong differential contacts. **a,** Schematic overview of determining differential contact count and differential contact hotspots. Hotspot regions were identified using a threshold of 28 differential contacts for Hap1 and Hap2 per window (within 10 Mb distance). **b,** Percentage of contacts identified as differential contacts per chromosome and haplotype. Notably, the parental genomes of H1 hESC are not known, thus haplotype labels do not correspond to a consistent parental origin across chromosomes. **c,** Fraction of imprinted and ASE genes expressed in H1, overlapping hotspot regions. Three different publicly available lists of imprinted gene sets are shown (Imprinted genes: red, Frost et al., 2011: blue, Kim et al., 2007: green). Numbers at the top of bars indicate total gene counts per group. **d,e,f,** Hotspot regions significantly overlap with at least one H3K27me3 (**d**) or H3K27ac (**e**) peaks or LAD region (**f**), compared with genome-wide expectation (permutation tests; p = 0.006, 0.050 and 0.137, respectively; 1000 permutations).

**Fig. S8:**
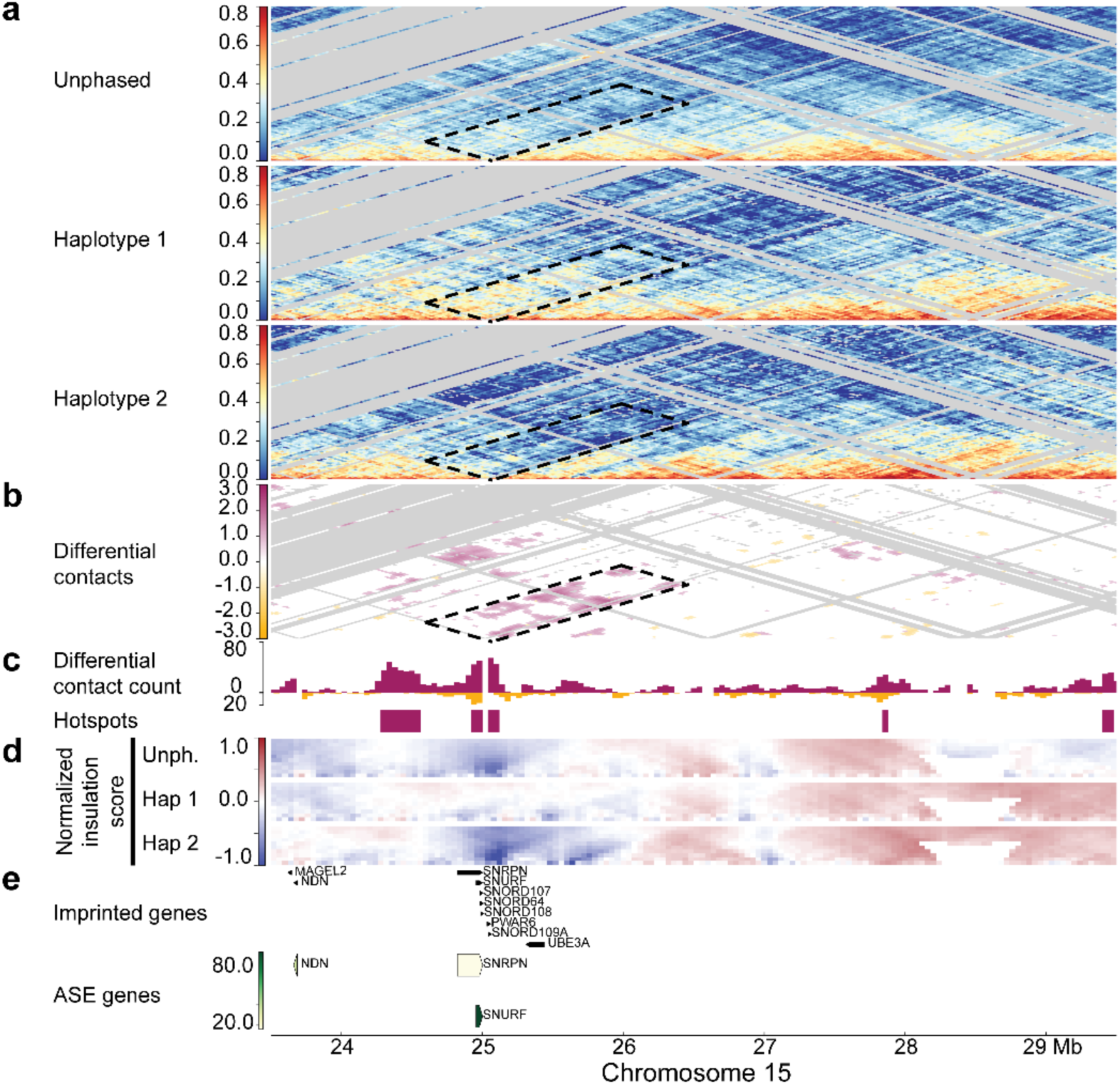
Cophased chromatin contact matrices from human GAM data reveal haplotype-specific differences within imprinted gene regions. CoPhasing-derived matrices in different regions of H1 hESC generated using 40 kb resolution and 10 Mb CoPhasing distance threshold. **a,** Unphased and cophased chromatin contact matrices in a 6 Mb region on chromosome 15 (chr15: 23.5 - 29.5 Mb). Red indicates frequent interactions between respective genomic regions and blue no interactions. Filtered regions in the matrices are depicted in light gray and regions with low mappability in GRCh38 genome are indicated below (mean mappability ≤ 0.1). **b,** Differential contacts as identified by permutation test. The directionality of the stronger contact is visualized by the color scale (red: stronger contact on haplotype 1, white: no difference detected, yellow: stronger contact on haplotype 2). **c,** Number of differential contacts per 40 kb bin based on permutation test results (Differential contact count) and below differential contact hotspot regions for both haplotypes. **d,** Normalized insulation score for unphased and cophased haplotype 1 and haplotype 2 respectively with insulation square sizes from 80 kb (bottom) to 800 kb (top). The color scale indicates local interaction frequency (red: higher local interaction frequency, blue: interaction drop). **e,** Expressed imprinted and ASE genes in H1 hESC with expression level of ASE genes indicated by the color scale filling the arrows (white: low expression, green: high expression).

**Fig. S9:**
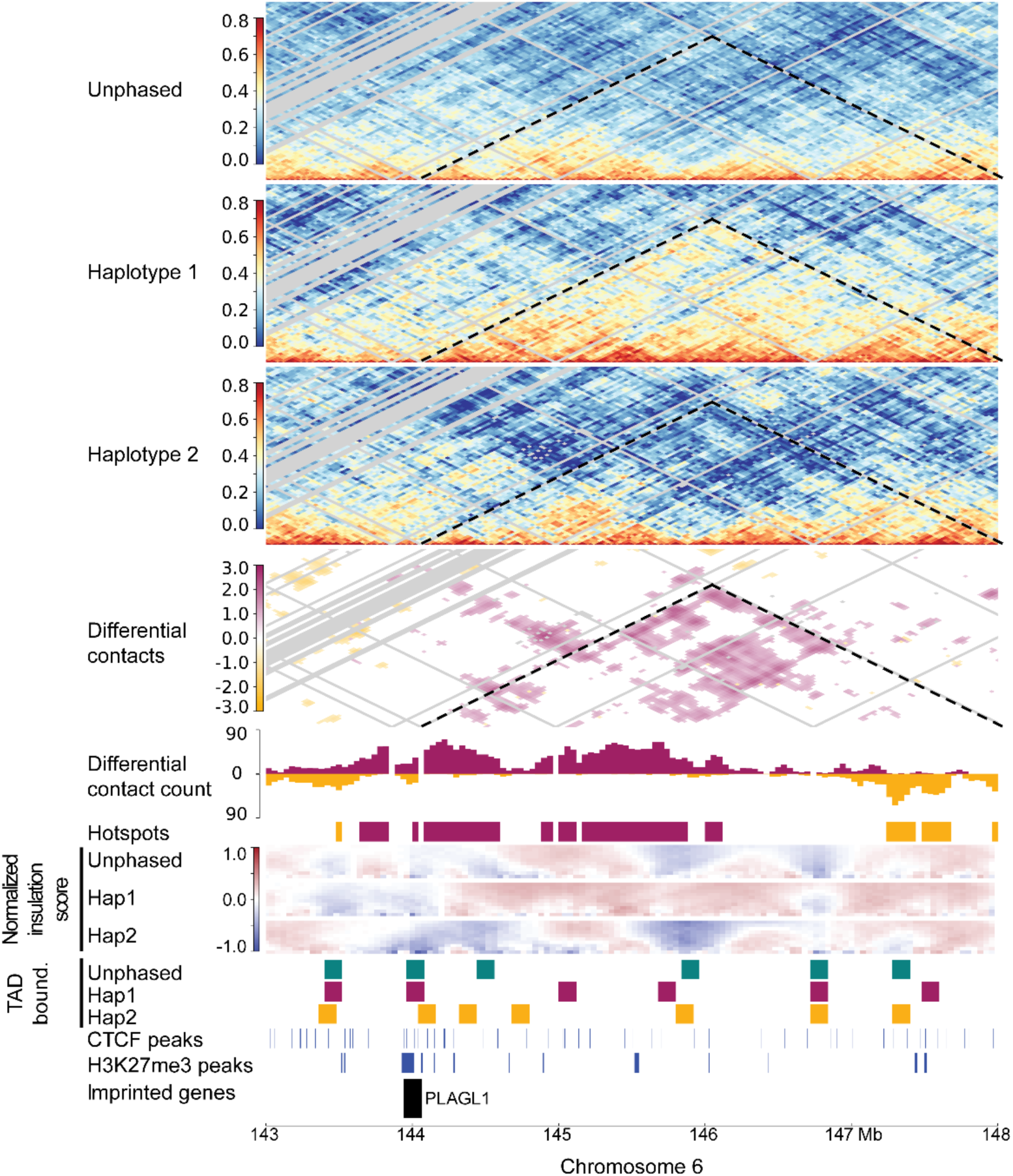
Region around imprinted gene *PLAGL1* shows haplotype-specific TAD boundaries. Haplotype-specific TAD boundaries underscoring structural variation in 3D genomic organization in 5 Mb region (chr 6: 143 - 148 Mb) around imprinted gene PLAGL1 (Frost et al., 2011). Unphased and cophased chromatin contact matrices and haplotype-specific chromatin contacts as identified by permutation test. Differential contact count and hotspot regions based on permutation test results are visualised per haplotype (Haplotype1: red, Haplotype2: yellow). Normalized insulation score for unphased and cophased haplotype 1 and haplotype 2 respectively with insulation square sizes from 80 kb (bottom) to 800 kb (top). The color scale indicates local interaction frequency (red: higher local interaction frequency, blue: interaction drop). TAD boundary location identified based on normalized insulation score using insulation square size of 400 kb for unphased and haplotype-specific data. Below, location of CTCF and H3K27me3 peaks. Location of the *PLAGL1* gene is depicted at the bottom.

**Fig. S10:**
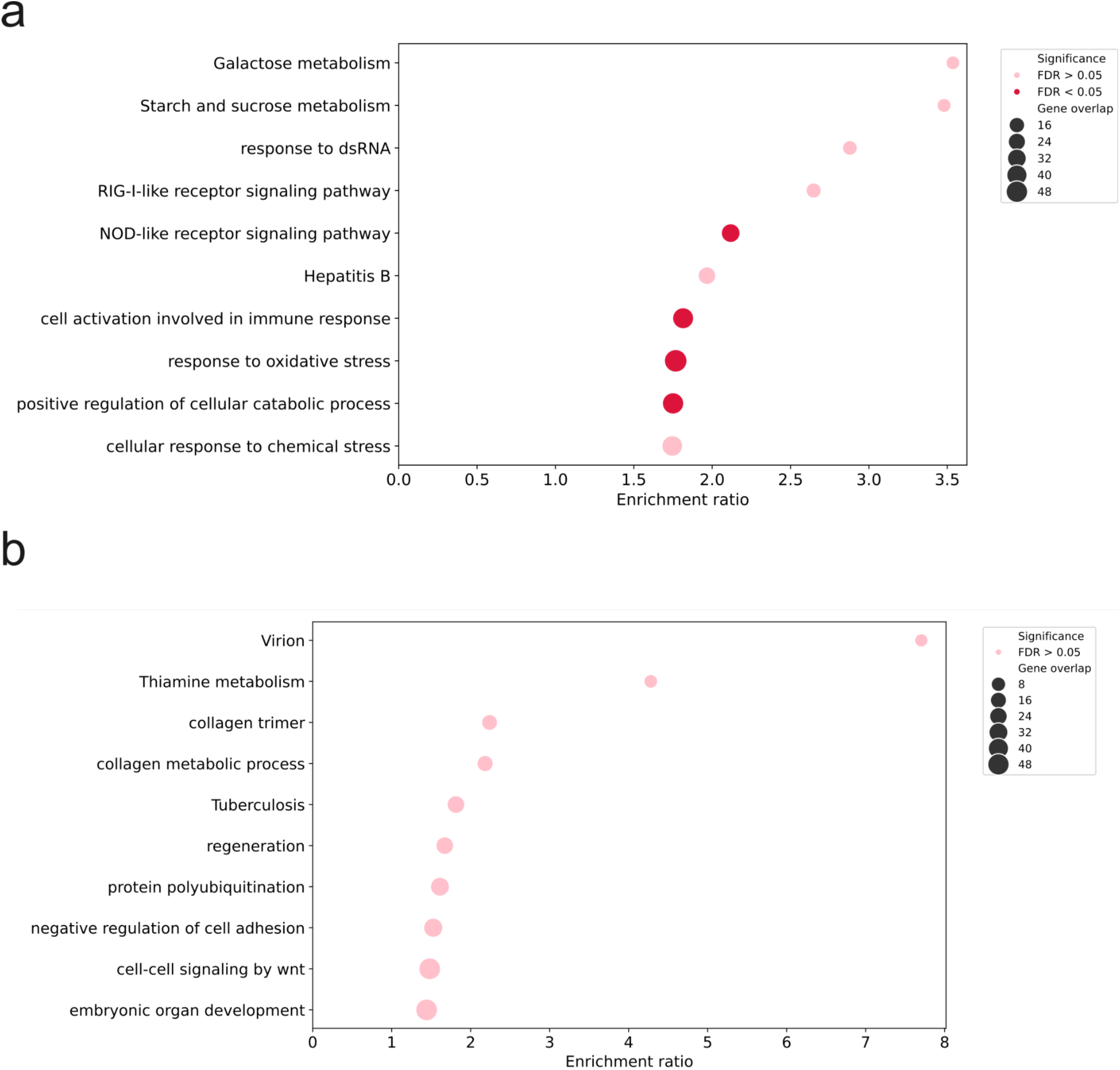
Gene Ontology (GO) enrichment analysis for genes found at **a**, common TAD boundaries and **b**, haplotype-specific TAD boundaries.

## Notes

https://doi.org/10.5281/zenodo.19740137

https://data.4dnucleome.org/experiments-seq/4DNEXVBJBGDP/

https://data.4dnucleome.org/experiments-seq/4DNEXL24V4SR/

## References

1. Kempfer, R. & Pombo, A. Methods for mapping 3D chromosome architecture. Nat. Rev. Genet. 21, 207–226 (2020).

2. Dekker, J. et al. The 4D nucleome project. Nature 549, 219–226 (2017).

3. Dekker, J. et al. An integrated view of the structure and function of the human 4D nucleome. Nature 649, 759–776 (2026).

4. Lupiáñez, D. G. et al. Disruptions of topological chromatin domains cause pathogenic rewiring of gene-enhancer interactions. Cell 161, 1012–1025 (2015).

5. Bendl, J. et al. The three-dimensional landscape of cortical chromatin accessibility in Alzheimer’s disease. Nat. Neurosci. 25, 1366–1378 (2022).

6. Girdhar, K. et al. Chromatin domain alterations linked to 3D genome organization in a large cohort of schizophrenia and bipolar disorder brains. Nat. Neurosci. 25, 474–483 (2022).

7. Gutnik, S., You, J. E., Sawh, A. N., Andriollo, A. & Mango, S. E. Multiplex DNA fluorescence in situ hybridization to analyze maternal vs. paternal C. elegans chromosomes. Genome Biol. 25, 71 (2024).

8. Dixon, J. R. et al. Chromatin architecture reorganization during stem cell differentiation. Nature 518, 331–336 (2015).

9. Richer, S. et al. Widespread allele-specific topological domains in the human genome are not confined to imprinted gene clusters. Genome Biol. 24, 40 (2023).

10. Irastorza-Azcarate, I. et al. Extensive folding variability between homologous chromosomes in mammalian cells. Mol. Syst. Biol. 21, 735–775 (2025).

11. Bae, B., Gu, K., Loftus, D. & Whipple, A. Allelic chromatin structure is a pervasive feature of imprinted domains and functions cooperatively with cis-acting long non-coding RNAs. Preprint at 10.64898/2025.12.02.691962 (2025).

12. Ijaz, J. et al. Haplotype-specific assembly of shattered chromosomes in esophageal adenocarcinomas. Cell Genomics 4, 100484 (2024).

13. Rao, S. S. P. et al. A 3D map of the human genome at kilobase resolution reveals principles of chromatin looping. Cell 159, 1665–1680 (2014).

14. The 1000 Genomes Project Consortium. A global reference for human genetic variation. Nature 526, 68–74 (2015).

15. Zhang, S. et al. DeepLoop robustly maps chromatin interactions from sparse allele-resolved or single-cell Hi-C data at kilobase resolution. Nat. Genet. 54, 1013–1025 (2022).

16. Servant, N. et al. HiC-Pro: an optimized and flexible pipeline for Hi-C data processing. Genome Biol. 16, 259 (2015).

17. Beagrie, R. A. et al. Complex multi-enhancer contacts captured by genome architecture mapping. Nature 543, 519–524 (2017).

18. Markowski, J. et al. GAMIBHEAR: whole-genome haplotype reconstruction from Genome Architecture Mapping data. Bioinformatics 37, 3128–3135 (2021).

19. Khalil, A. et al. Chromosome territories have a highly nonspherical morphology and nonrandom positioning. Chromosome Res. Int. J. Mol. Supramol. Evol. Asp. Chromosome Biol. 15, 899–916 (2007).

20. Beagrie, R. A. et al. Multiplex-GAM: genome-wide identification of chromatin contacts yields insights overlooked by Hi-C. Nat. Methods 20, 1037–1047 (2023).

21. Markowski, J., et al. Extended data for manuscript ‘Haplotype-resolved genome architecture mapping uncovers pervasive structural heterogeneity between human homologous chromosomes’. Zenodo 10.5281/zenodo.19740137 (2026).

22. Kumar, P. et al. Nucleolus and centromere Tyramide Signal Amplification-Seq reveals variable localization of heterochromatin in different cell types. *Commun*. Biol. 7, 1135 (2024).

23. Frost, J. M. et al. The effects of culture on genomic imprinting profiles in human embryonic and fetal mesenchymal stem cells. Epigenetics 6, 52–62 (2011).

24. Kim, K.-P. et al. Gene-specific vulnerability to imprinting variability in human embryonic stem cell lines. Genome Res. 17, 1731–1742 (2007).

25. Crane, E. et al. Condensin-driven remodelling of X chromosome topology during dosage compensation. Nature 523, 240–244 (2015).

26. Winick-Ng, W. et al. Cell-type specialization is encoded by specific chromatin topologies. Nature 599, 684–691 (2021).

27. Akgol Oksuz, B., et al. Systematic evaluation of chromosome conformation capture assays. Nat. Methods 18, 1046–1055 (2021).

28. Dixon, J. R. et al. Topological domains in mammalian genomes identified by analysis of chromatin interactions. Nature 485, 376–380 (2012).

29. Özçelik, T. et al. Small nuclear ribonucleoprotein polypeptide N (SNRPN), an expressed gene in the Prader-Willi syndrome critical region. Nat. Genet. 2, 265–269 (1992).

30. Arnoult, N. et al. Regulation of DNA Repair pathway choice in S/G2 by the NHEJ inhibitor CYREN. Nature 549, 548–552 (2017).

31. Mariner, B. L. et al. Diverse Patterns of Allele-Specific Expression in Healthy Human Tissues. Preprint at 10.1101/2025.10.14.682127 (2025).

32. Bar, S. & Benvenisty, N. Epigenetic aberrations in human pluripotent stem cells. EMBOJ. 38, EMBJ2018101033 (2019).

33. Gribnau, J., Hochedlinger, K., Hata, K., Li, E. & Jaenisch, R. Asynchronous replication timing of imprinted loci is independent of DNA methylation, but consistent with differential subnuclear localization. Genes Dev. 17, 759–773 (2003).

34. Thomson, J. A. et al. Embryonic Stem Cell Lines Derived from Human Blastocysts. Science 282, 1145–1147 (1998).

35. Reiff, S. B. et al. The 4D Nucleome Data Portal as a resource for searching and visualizing curated nucleomics data. Nat. Commun. 13, 2365 (2022).

36. Hinrichs, A. S. et al. The UCSC Genome Browser Database: update 2006. Nucleic Acids Res. 34, D590–D598 (2006).

37. Langmead, B. & Salzberg, S. L. Fast gapped-read alignment with Bowtie 2. Nat. Methods 9, 357–359 (2012).

38. Li, Z. et al. RGT: a toolbox for the integrative analysis of high throughput regulatory genomics data. BMC Bioinformatics 24, 79 (2023).

39. Marco-Sola, S., Sammeth, M., Guigó, R. & Ribeca, P. The GEM mapper: Fast, accurate and versatile alignment by filtration. Nat. Methods 9, 1185–1188 (2012).

40. ENCODE Project Consortium. An integrated encyclopedia of DNA elements in the human genome. Nature 489, 57–74 (2012).

41. Mei, S. et al. Cistrome Data Browser: a data portal for ChIP-Seq and chromatin accessibility data in human and mouse. Nucleic Acids Res. 45, D658–D662 (2017).

42. Ji, X. et al. 3D Chromosome Regulatory Landscape of Human Pluripotent Cells. Cell Stem Cell 18, 262–275 (2016).

